# The membrane distal domain of CD16a allosterically regulates NK cell ADCC

**DOI:** 10.64898/2026.07.03.736431

**Authors:** Tania Cid, Monica Fernandez-Quintero, Hijab Fatima, Ema Robinson, Braden Christenson, Johannes Loeffler, Daniel Leaman, Ryan Lin, Kayla Xu, Jessica Matthias, Scott C. Henderson, Kathryn Spencer, Joseph Jardine, Michael B. Zwick, Andrew B. Ward, Emily M. Mace, Charles Daniel Murin

## Abstract

Antibody-dependent cellular cytotoxicity (ADCC) by natural killer (NK) cells is mediated by the activating IgG receptor CD16a (FcγRIIIa), yet the molecular mechanisms governing receptor activation remain poorly understood. We demonstrate that the membrane-distal domain 1 (D1) of CD16a functions as an allosteric checkpoint that controls ADCC independently of IgG-Fc binding. A nanobody, C28, that binds an electronegative patch in D1 dose-dependently blocks NK cell ADCC against multiple therapeutic antibodies without affecting direct cytotoxicity. A second nanobody, C21, binding an adjacent D1 epitope has no such effect. Cryo-EM structures of the CD16a-IgG-nanobody complex reveal that C28 allosterically competes with core-fucosylated IgG and stabilizes a closed D1 conformation resembling unliganded receptor, even when Fc is bound. Molecular dynamics simulations show that occupation of the D1 epitope rigidifies the IgG-binding site, stabilizing CD16a overall in contrast with IgG binding alone. The nanobody C28 restricts CD3ζ phosphorylation in both resting and ADCC-activated NK cells, revealing tonic inhibitory control upstream of the signaling cascade. Using MINFLUX nanoscopy, we also show that CD16a forms dimers of ∼9 nm spacing on the NK cell surface, a geometry unaltered by the ADCC-enhancing L48H polymorphism. Drawing on structural parallels with the IgE receptor FcεRI, which is held inactive as a cholesterol-stabilized dimer, we propose that CD16a dimerization through D1 contacts represents a conserved autoinhibitory mechanism among Fc receptors. Consistent with this model, structure-guided disruption of the C28 epitope in NK-92 cells enhances ADCC potency and killing kinetics, providing a blueprint for engineering improved cellular immunotherapeutics.

## Introduction

Natural killer (NK) cells are central effectors of antibody-based cancer immunotherapies, mediating antibody-dependent cellular cytotoxicity (ADCC) almost exclusively through the activating IgG receptor FcγRIIIa (CD16a). Monoclonal antibodies such as Rituximab and Trastuzumab leverage this axis to recruit NK cells against CD20-positive lymphoma and HER2-positive cancers, respectively, and optimizing ADCC is a primary goal in both therapeutic antibody design and NK cell engineering. Despite this clinical importance, a coherent mechanistic framework for how CD16a activation is regulated, and how it might be rationally enhanced, has remained elusive. The immune synapse (IS) formed between an NK cell and its target has been characterized by super-resolution imaging^1–4^, CD16a clustering^5–7^, and CD3ζ phosphorylation^8–11^. Mechanotransduction^12^ inputs have each been implicated in ADCC, but how these elements are integrated at the receptor level is unclear. We recently demonstrated that CD16a constitutively forms pairs on the NK cell surface that do not undergo extensive clustering upon ADCC activation^13^, implying that receptor oligomeric organization plays a regulatory role beyond simple ligand engagement.

CD16a is a single-pass transmembrane glycoprotein whose ectodomain comprises two Ig-like folds: a membrane-proximal domain 2 (D2) that directly engages IgG-Fc, and a membrane-distal domain 1 (D1) whose function is poorly defined. The 3G8 antibody blocks IgG binding through the D2 epitope^14^, while the B73.1 antibody paradoxically inhibits ADCC without displacing IgG, targeting D1 by an unexplained mechanism^15^. A naturally occurring D1 substitution at residue 48 (L48H/R) eliminates association of CD2 on the NK cell surface and reduces direct cytotoxicity yet enhances ADCC by compacting the IS and promoting serial killing^16–18^. These observations collectively point to D1 as a functional regulatory element, but the structural basis and mechanism of its role in ADCC have not been established.

Two single-domain antibodies (nanobodies), C21 and C28, were previously identified as binding CD16a D1 at non-overlapping epitopes outside the IgG-Fc binding site^19^. Using these nanobodies as molecular probes, we unexpectedly found that C28, but not C21, dose-dependently blocks ADCC across multiple therapeutic antibodies without affecting direct cytotoxicity. A cryo-EM structure of CD16a in complex with IgG Fc, Fab P2C-47, and both nanobodies reveals the structural basis for this distinction and defines an allosteric regulatory site in D1. Biolayer interferometry, stimulated emission depletion (STED)-based CD3ζ imaging, MINFLUX nanoscopy, and molecular dynamics simulations together demonstrate that this site controls receptor conformation and downstream signaling. Structural parallels with the recently determined inactive-dimer structure of the IgE receptor FcεRI^20^ support a conserved autoinhibitory mechanism across the Fc receptor family. Finally, structure-guided mutagenesis of the C28 epitope in NK-92 cells enhances ADCC potency and killing kinetics, providing a blueprint for engineering NK cells with improved therapeutic activity.

## Results

### A nanobody to CD16a blocks NK cell ADCC

Two single domain antibodies (also known as nanobodies), were previously generated through vaccination of a llama with the human CD16a ectodomain^19^. The nanobodies C21 and C28 were characterized as having non-overlapping epitopes that bind distal to the 3G8 antibody epitope, which itself overlaps with the IgG-Fc binding site in D2 of CD16a. This indicates that these nanobodies do not directly block IgG binding to CD16a. Fortuitously, we first tested that these antibodies did not inhibit NK cell ADCC before utilizing them for single molecule light microscopy (SMLM) studies to analyze CD16a distribution during NK cell ADCC. Using our previously established impedance assay^13^, we first measured the dose response ADCC of NK-92^CD16a^ cells with an endogenously activated CD16a gene^21^ against SKOV3 cells coated with Trastuzumab (Fig. 1A). We next treated SKOV3 cells with a high concentration of Trastuzumab where 100% of targets are killed, dosed in either C21 or C28 nanobody, and measured real-time killing kinetics. While C21 did not alter cytotoxicity, C28 dose-dependently blocked ADCC with an IC_50_ of ∼1 µg/mL (Fig. 1B). We also observed C28 blocking of ADCC using antibodies targeting a different HER2 epitope (Pertuzumab) and different protein target (Cetuximab, EGFR) (Fig. 1C, Fig. S1A), though with differing IC_50_ values, possibly indicating that target proteins and epitopes influence ADCC activity. Primary NK cells derived from PBMCs of healthy donors were also blocked from performing ADCC in the presence of C28 (Fig. 1D).

**Figure 1.**
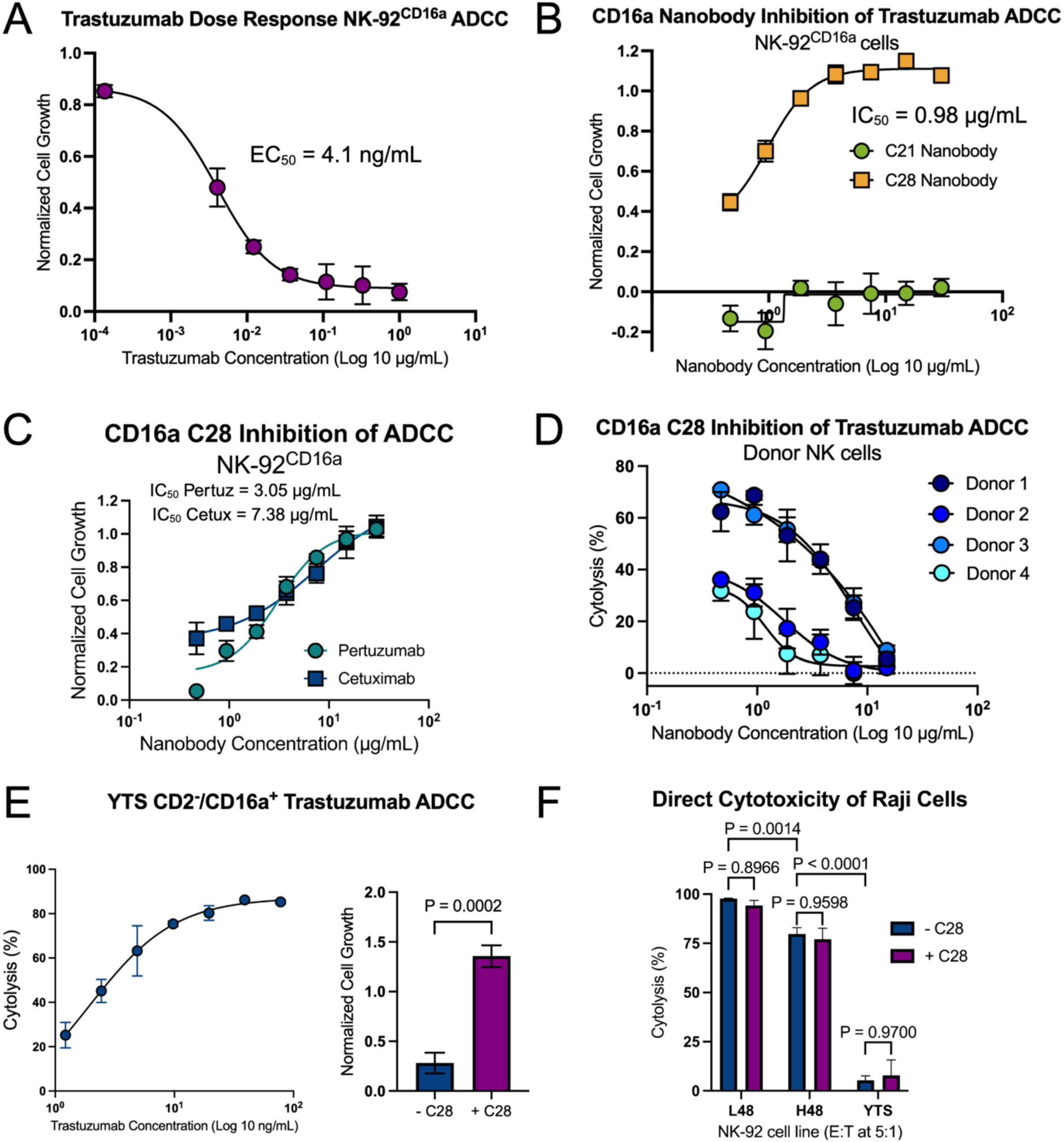
Nanobody C28 blocks NK cell ADCC activity but not direct cytotoxicity. A) Dose-response ADCC activity of NK-92/CD16a cells using Trastuzumab to target SKOV3 cells (E:T ratio of 5:1). B) Dose-response inhibition by C28 but not C21 of Trastuzumab mediated ADCC by NK-92/CD16a cells, using a concentration of Trastuzumab (78 ng/mL) that induces complete cytolysis of SKOV3 cells (E:T ratio of 5:1). C) C28 dose-dependently blocks the NK-92/CD16a mediated ADCC activity of Pertuzumab (anti-HER2) and Cetuximab (anti-EGFR) (E:T ratio of 2:1). D) C28 inhibits primary NK cell-mediated ADCC of Trastuzumab toward SKOV3 cells, though some donor-to-donor variability is observed. E) YTS NK cells lack CD2 and poorly perform direct cytotoxicity but are CD16a positive and can perform ADCC normally (Trastuzumab dose-response toward SKOV3 cells shown on the left). Shown on the right, YTS NK cells are blocked from performing ADCC in the presence of C28 (Trastuzumab at 78 ng/mL, C28 at 30 µg/mL). A Welch’s t-test was performed to determine significance. F) NK-92/CD16a with the WT residue at L48 or the H48 variant, along with YTS NK cells, were tested for their ability to directly kill Raji cells in the absence or presence of C28 (E:T ratio of 2:1). At the same E:T ratio, The CD16a H48 variant performs worse than the WT L48 variant, while YTS cells have little to no cytotoxicity. In all cases, C28 does not alter natural cytotoxicity. Statistical significance was determine using an ordinary one-way ANOVA test. EC_50_ is the effective concentration at 50% cytotoxicity. IC_50_ is the inhibitory concentration at 50% antibody inhibition.

We next wondered if C28 binding interferes with CD2’s interaction with CD16a. In NK cells bearing a substitution at residue 48 (L48H/R), CD2 interaction is lost and ADCC is significantly enhanced^16,22^. We tested the effects of C28 on YTS^CD16a^ NK cells, which do not express CD2, and found that ADCC activity was also blocked (Fig. 1E). The addition of the CD2 ectodomain (eCD2) (Fig. S1B) or the addition of an anti-CD2 antibody (Fig. S1C) did not affect ADCC activity. We next treated NK-92/CD16^H48^ cells with C28 and saw a similar, though to a lesser degree, loss in ADCC activity compared to NK-92/CD16^L48^ (Fig. S1D-G). Finally, we saw no decrease in direct cytotoxicity in the presence of C28 (Fig. 1F). These data collectively suggested that disruption of cytotoxicity via the binding of C28 was specific for ADCC and independent of CD2 and mechanisms of direct cytotoxicity. Further, these data show that the C28 epitope on CD16a D1 is specifically responsible for this effect.

### C28 but not C21 competes with fucosylated but not afucosylated IgG for binding to CD16a

To determine if C28 competes with IgG for binding, we employed a series of competition studies using biolayer interferometry (BLI) (Fig. 2A). As expected, C21 and IgG did not compete for binding in either assay (Fig. 2B-C). However, the presence of C28 actively competed for Trastuzumab binding (Fig. 2B). In the reverse assay, there was some IgG binding, but it was markedly reduced (Fig. 2C). Notably, the commercial antibodies used were made in CHO cells and presumably contain fucose at the core N297 glycan. Therefore, we questioned if an afucosylated IgG could bind in the presence of C28, so we compared binding of the anti-HIV Env antibody PGT-121 prepared in-house either with or without core fucose. As with Trastuzumab, the fucosylated version was actively outcompeted by C28 (Fig. 2D-E). However, the afucosyalted version was able to recover some binding after being displaced (Fig. 2F) and bound well to CD16a if C28 was already present, though less than when C21 was present (Fig. 2G). Crucially, the CD16a we used in these assays is fully glycosylated. Likely, trimming glycans or preventing processing would enhance IgG binding even further, as has been previously shown^23,24^. By combining these observations with previous epitope mapping, we hypothesized that C28 may be allosterically competing with IgG binding rather than sterically.

**Figure 2.**
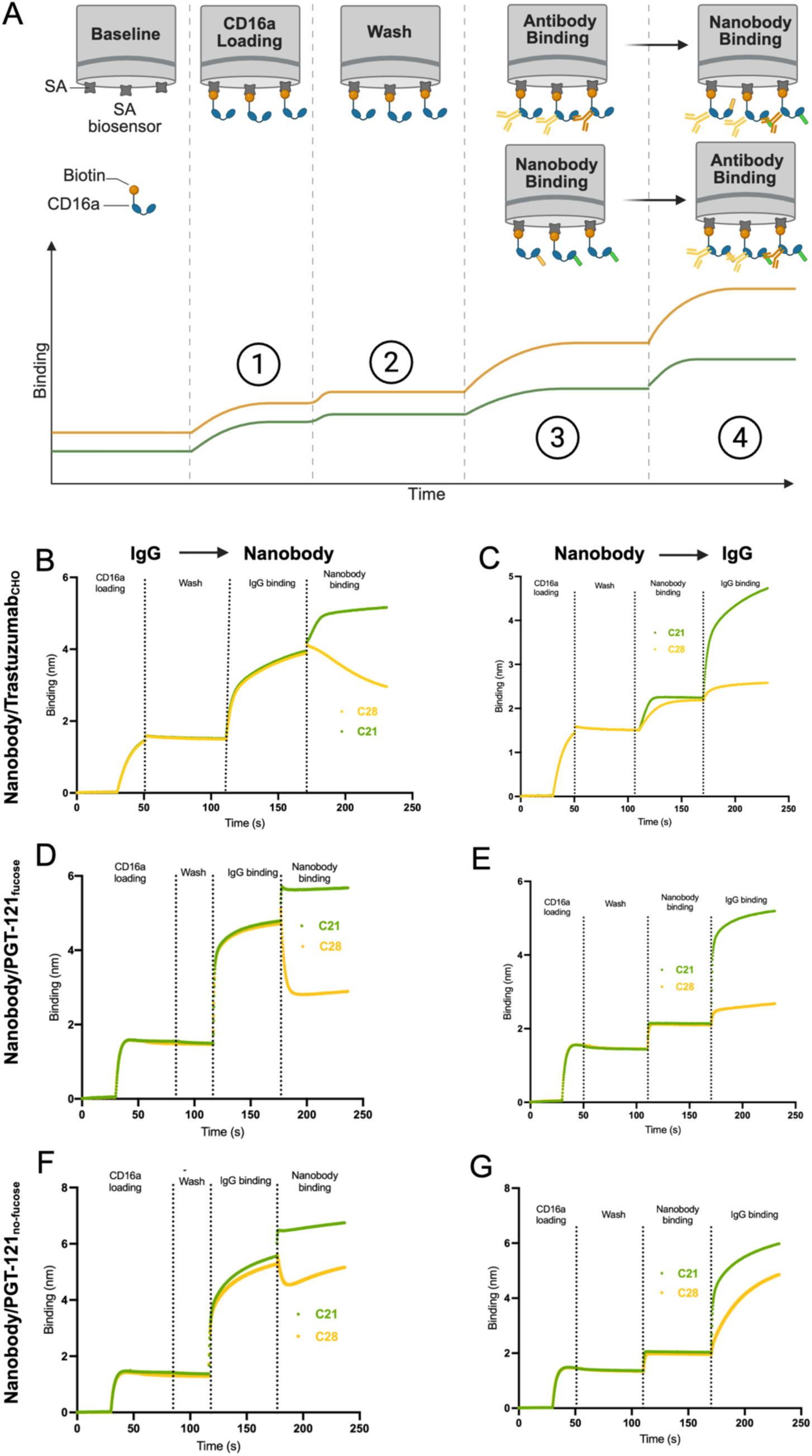
Fucosylated but not fucose-free IgG is blocked from binding in the presence of C28 nanobody. A) Schematic showing the experimental setup for all competition assays using biolayer interferometry (BLI): 1. Streptavidin (SA) biosensors are first activated in kinetics buffer followed by loading of biotinylated CD16a ectodomains (via their C-terminal domain). 2. Biosensors are then washed in kinetics buffer. 3. Next, either nanobody or IgG is first bound to the biosensors. 4. Immune complex coated biosensors are dipped into the opposite ligand, which contains the same concentration of the previous ligand. B) CHO cell produced Trastuzumab was first bound to biosensors, followed by nanobody, indicating C28 causes the IgG to dissociate while C21 does not. C) Opposite to part B, showing very little IgG binding in the presence of C28 compared to C21. D) CHO cell produced PGT-121 (containing fucose) rapidly dissociates from CD16a in the presence of C28. E) Opposite to part D, very little fucosylated PGT-121 binds to CD16a in the presence of C28 compared to C21. F) CHO cell produced PGT-121 lacking fucose (afucosylated) dissociates in the presence of C28 but binding is quickly recovered. G) Opposite to part F, fucose-free PGT-121 readily binds to CD16a in the presence of C28.

Notably, our binding assays are performed with the monomeric, extracellular CD16a domain, lacking the cellular context where we have shown CD16a to be dimeric. Further, CD16a likely has additional *cis-*binding partners, such as CD2 for example, that may influence activation. We therefore compared the ADCC activity of core fucosylated versus afucosylated Cetuximab using NK-92/CD16a^L48^ cells and determined how C28 differentially affects this activity. As expected, C28 dose-dependently blocked ADCC with CHO-cell produced Trastuzumab, as we showed before (Fig. S2). Though the potency of fucose-free Cetuximab is higher than fucosylated at the same concentration, ADCC was also attenuated in the presence of C28 and to a relatively higher degree (Fig. S2). This is despite demonstration that fucose-free IgG is competent to bind CD16a monomers in the presence of C28. This suggests that C28 either prevents dimer to monomer transition on the cell surface, prevents an IgG-induced conformational change that is necessary for ADCC-based activation, and/or mimics the binding of an unknown *cis*-acting protein ligand.

### CD16a nanobodies bind to non-overlapping epitopes in D1 distinct from the IgG binding site

We next sought to determine the specific binding sites of C21 and C28 and so made a complex of proteins expressed and purified from mammalian cells including CD16a ectodomain bound to IgG Fc, Fab P2C-47 (as a fiducial that does not block IgG binding)^25^, and nanobodies C21 and C28. To increase the binding affinity and stability of our complex, we treated cells expressing IgG Fc with 2-deoxy-2-fluor-fucose to remove core fucose from our recombinant Fc. We also used a construct of CD16a ectodomain missing glycans at N38, N74 and N169^26,27^, which are not essential for IgG binding or ADCC activity, and treated cells with kifunensine to simplify glycans on CD16a to increase IgG Fc binding. Using cryo-electron microscopy (cryo-EM), we solved a structure of the full complex to a global resolution of 4.2 Å (Fig. 3A, Fig. S3).

**Figure 3.**
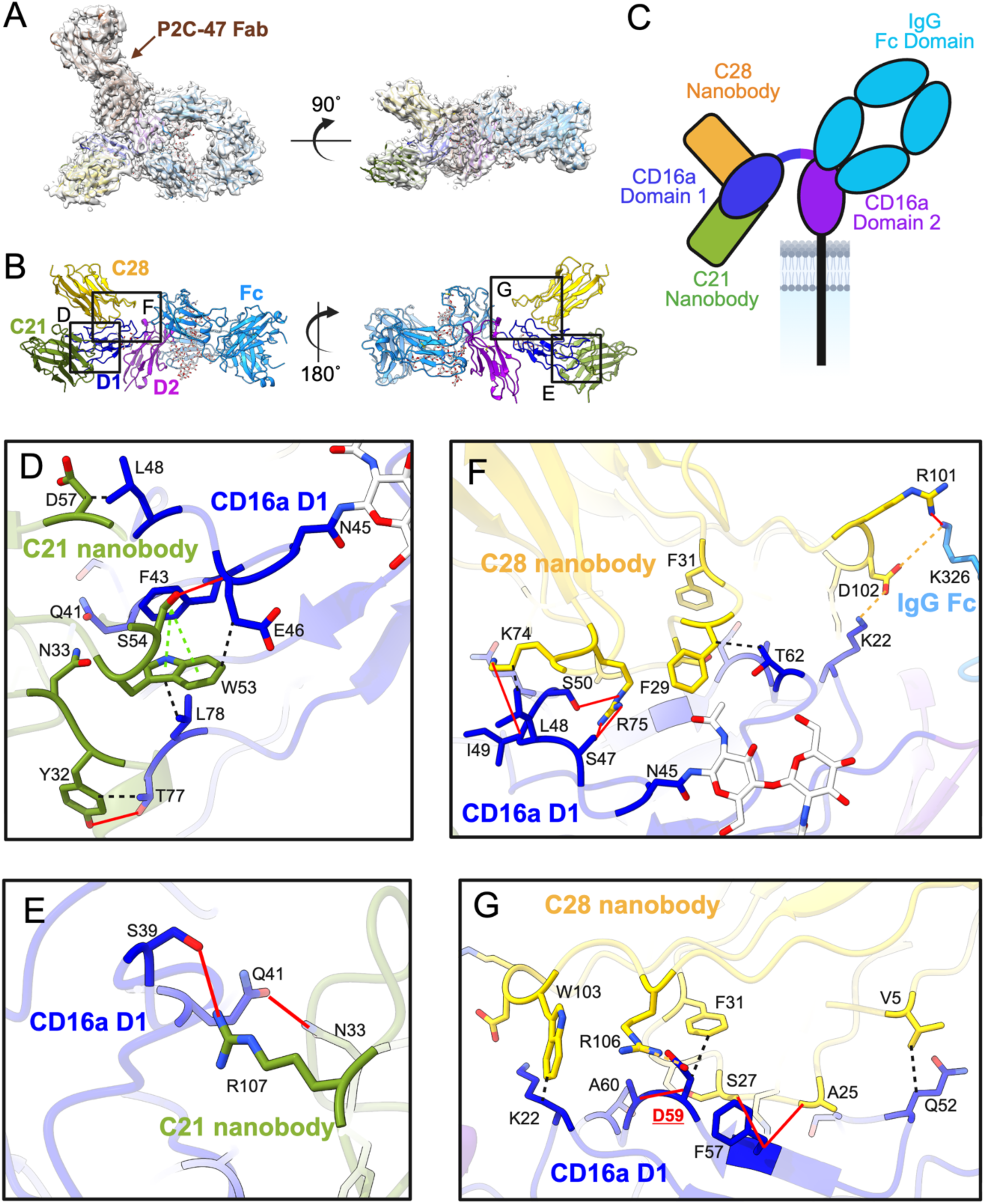
Cryo-EM structure of CD16a bound to IgG Fc, Fab P2C-47, and nanobodies C21 and C28. A) Final map and model of the EM complex showing top (left) and side (right) views, with fiducial Fab P2C-47 indicated. B) Overall model of the EM complex with components and CD16a subdomains indicated. Boxed areas are of zoomed in epitope/paratope details below. C) Cartoon schematic of the overall EM complex to indicate the approximate location of all ligands in relation to subdomains on CD16a. D-G) Details of the C28 and C21 epitopes from regions indicated in part B. All residues that contribute to contacts between CD16a, Fc and nanobodies are shown, including the type of interaction. Solid red lines indicate hydrogen bonds, dotted black lines indicate hydrophobic interactions, dotted green lines indicate pi stacking, and dotted yellow lines indicate salt bridges. Residues indicated in red are among those identified as being important for binding.

We could clearly observe that C21 and C28 bind to non-overlapping epitopes in D1of CD16a, which do not overlap with the IgG binding site (Fig. 3B-C). C21 binds to the tip of D1 while C28 binds to the top of D1 (Fig. 3C. Fig. S4). Using our model, we performed protein-protein interaction analysis using Protein-Ligand Interaction Profiler (PLIP), generating a list of primary interactions for both nanobodies (Table 1). Though resolution was lower for C21 density, we could confidently dock and resolve its structure, showing that C21 binds exclusively to D1, though more on the side. Primary interactions on CD16a include hydrophobic interactions with D46, L48, T77, and L78, pi-stacking with F43, and hydrogen bonds with S39, Q41, E46, and T77 (Table 1, Fig. 3D-E).

**Table 1.** Protein-protein interactions of nanobodies with CD16a and IgG Fc. To characterize the epitope-paratope interactions of nanobodies C21 and C28, the final refined model of the cryo-EM complex was submitted to PLIP analysis. Listed are the interacting residues, their distance, and the type of interaction. These interactions are visually represented in Figure 4.

| Protein A | Residue A | Protein B | Residue B | Distance (Å) | Type |
| --- | --- | --- | --- | --- | --- |
| CD16a | K22 | C28 | W103 | 3.97 | Hydrophobic |
| CD16a | L48 | C28 | K74 | 3.72 | Hydrophobic |
| CD16a | Q52 | C28 | V5 | 3.87 | Hydrophobic |
| CD16a | D59 | C28 | F31 | 3.84 | Hydrophobic |
| CD16a | T62 | C28 | F29 | 3.69 | Hydrophobic |
| CD16a | S47 | C28 | R75 | 2.65 | H-bond |
| CD16a | S47 | C28 | R75 | 2.49 | H-bond |
| CD16a | I49 | C28 | K74 | 2.59 | H-bond |
| CD16a | S50 | C28 | R75 | 2.38 | H-bond |
| CD16a | F57 | C28 | A25 | 3.09 | H-bond |
| CD16a | F57 | C28 | S27 | 2.71 | H-bond |
| CD16a | A60 | C28 | S27 | 2.52 | H-bond |
| CD16a | K22 | C28 | D102 | 3.94 | Salt bridge |
| CD16a | D59 | C28 | R106 | 4.63 | Salt bridge |
| IgG Fc | K326 | C28 | R101 | 2.61 | H-bond |
| IgG Fc | K326 | C28 | D102 | 5.48 | Salt bridge |
| CD16a | E46 | C21 | W53 | 3.42 | Hydrophobic |
| CD16a | L48 | C21 | D57 | 3.55 | Hydrophobic |
| CD16a | T77 | C21 | Y32 | 3.37 | Hydrophobic |
| CD16a | L78 | C21 | W53 | 3.67 | Hydrophobic |
| CD16a | S39 | C21 | R107 | 2.93 | H-bond |
| CD16a | Q41 | C21 | N33 | 2.41 | H-bond |
| CD16a | E46 | C21 | S54 | 2.32 | H-bond |
| CD16a | T77 | C21 | Y32 | 3.46 | H-bond |
| CD16a | F43 | C21 | W53 | 5.47 | Pi-stacking |
| CD16a | F43 | C21 | W53 | 4.49 | Pi-stacking |

The CDR3 of C28 reaches across the top of CD16a and has a single interaction with IgG Fc at K326, characterized by a salt bridge and hydrogen bond (Fig. 3F). The remainder of the C28 epitope is contained to the top portion of CD16a, forming interactions along a peptide from residues 47-62 and an additional salt bridge with K22 (Fig. 3F-G, Table 1). Notably, the epitope where C28 binds overlaps with a patch of highly electronegative residues (Fig. S5A). Collectively, our structures demonstrate that C28 binds to CD16a D1 in an epitope distinct from the IgG-Fc binding domain and from that of C21 (which does not alter NK cell ADCC). The epitope of C28 therefore likely defines a key site for regulating NK cell ADCC through the membrane distal D1.

### CD16a nanobody C28 alters immune synapse structure and CD3-zeta phosphorylation

We next wanted to determine at what level of signaling CD16a D1 regulates NK cell ADCC. We previously have used phosphorylated CD3-zeta (pCD3ζ) as a proxy for ADCC activated NK cells by measuring pCD3ζ with confocal microscopy^13^. To more arcuately describe pCD3ζ foci, here we used STED microscopy, which provides a 10-fold increase in spatial resolution over diffraction-limited confocal microscopy. NK-92/CD16a^L48^ cells were used to create immune synapses on supported lipid bilayers (SLBs) coated in HER2 and ICAM either with or without Trastuzumab and with or without C28. After fixing and labeling cells, we determined the plane of the synapse using phalloidin-488 counterstain and imaged STAR-Red labeled pCD3ζ foci, subsequently measuring foci density and number.

We found that the phosphorylation of CD3ζ is significantly decreased in cells treated with C28 (Fig. 4A-B). We routinely observe basal levels pCD3ζ in resting NK cells, however even resting pCD3ζ levels were lower in the presence of C28 without Trastuzuamb (Fig. 4A). To confirm our observations, we also attached primary donor NK cells to glass coated with ICAM ± Rituximab. In agreement with our NK-92 data on SLBs, we similarly observed that C28 disrupts CD3ζ phosphorylation in both resting and antibody-activated cells (Fig. 4C-D). These data indicate that blocking the C28 epitope in CD16a D1 alters downstream phosphorylation of the intracellular immune tyrosine activating motifs (ITAMs) on CD3ζ, a co-receptor associated with CD16a that is essential for signaling cascades responsible for ADCC based cytotoxicity.

**Figure 4.**
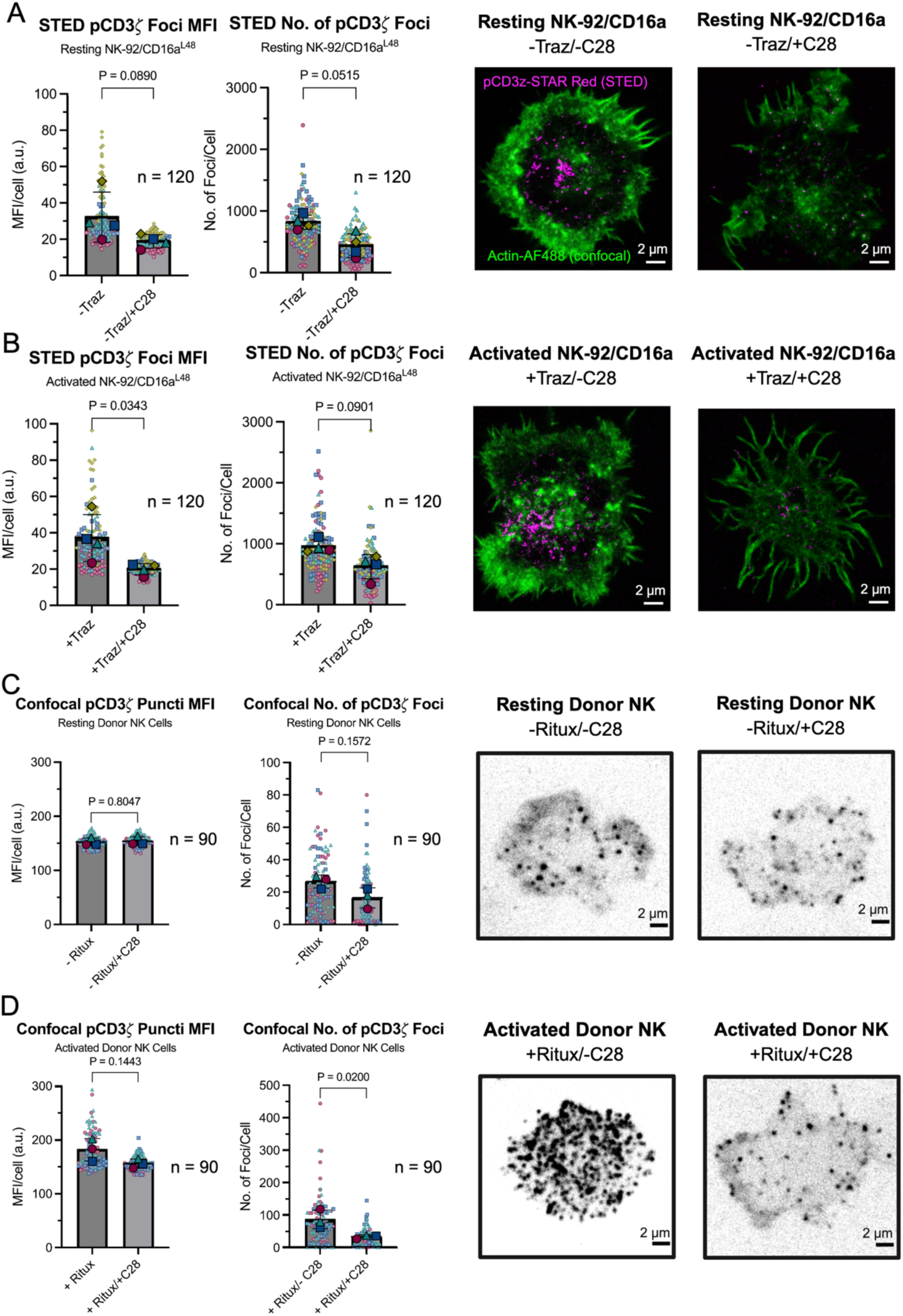
C28 nanobody binding to CD16a D1 alters phosphorylation of CD3ζ in resting and activated NK cells. A) NK-92/CD16a cells were coated on supported lipid bilayers (SLBs) containing ICAM and HER2, with and without C28 (30 µg/mL), and allowed to bind for 10 min before fixing and permeabilizing. Cells were stained and phosphor- CD3ζ (pCD3ζ) mean fluorescence intensity (MFI) and the number of pCD3ζ puncti (middle) were quantified using STED imaging. B) Similar to part A, cells were coated onto SLBs in the presence of Trastuzumab (10 µg/mL), with or without C28, and pCD3ζ puncti were similarly quantified from STED data. C) Primary NK cells were plated on ICAM-1 (10µg/mL) for 25 minutes to allow for synapse formation before fixing and permeabilizing. Cells were stained with pCD3ζ and the number of foci and MFI was quantified. D) Primary NK cells were plated on glass containing ICAM-1 + Rituximab (10µg/mL) for 25 minutes to allow for synapse formation before fixing and permeabilizing. Cells were stained for pCD3ζ and the number of foci and MFI was quantified. All data are represented using SuperPlots^67^. Three to four biological replicates were performed (n indicated, small symbols) and the mean was calculated (large symbols). Statistical significance was determined using un-paired, lognormal Welch’s t-tests, comparing means of each biological replicate. a.u. = arbitrary unit.

We also observed total CD3ζ at the immune synapse to gain more insight into the organization of the receptor itself. Using the same experimental setup for the STED analysis above, we instead labeled CD3ζ with the clone 6B10.2, that recognizes both the unphosphorylated and phosphorylated states (intracellular) and again measured CD3ζ foci density and number. In contrast to phosphorylated CD3ζ, we did not detect any significant differences in total CD3ζ at the synapse (Fig. 5A-B). Indeed, total CD3ζ in cells is consistent across all conditions when measured by flow cytometry (Fig. S6), indicating that the decrease of CD3ζ phosphorylation in the presence of C28 that we observed is most likely due to a disruption of phosphorylation directly, presumably by restricting kinase access, rather than changes in total CD3ζ.

**Figure 5.**
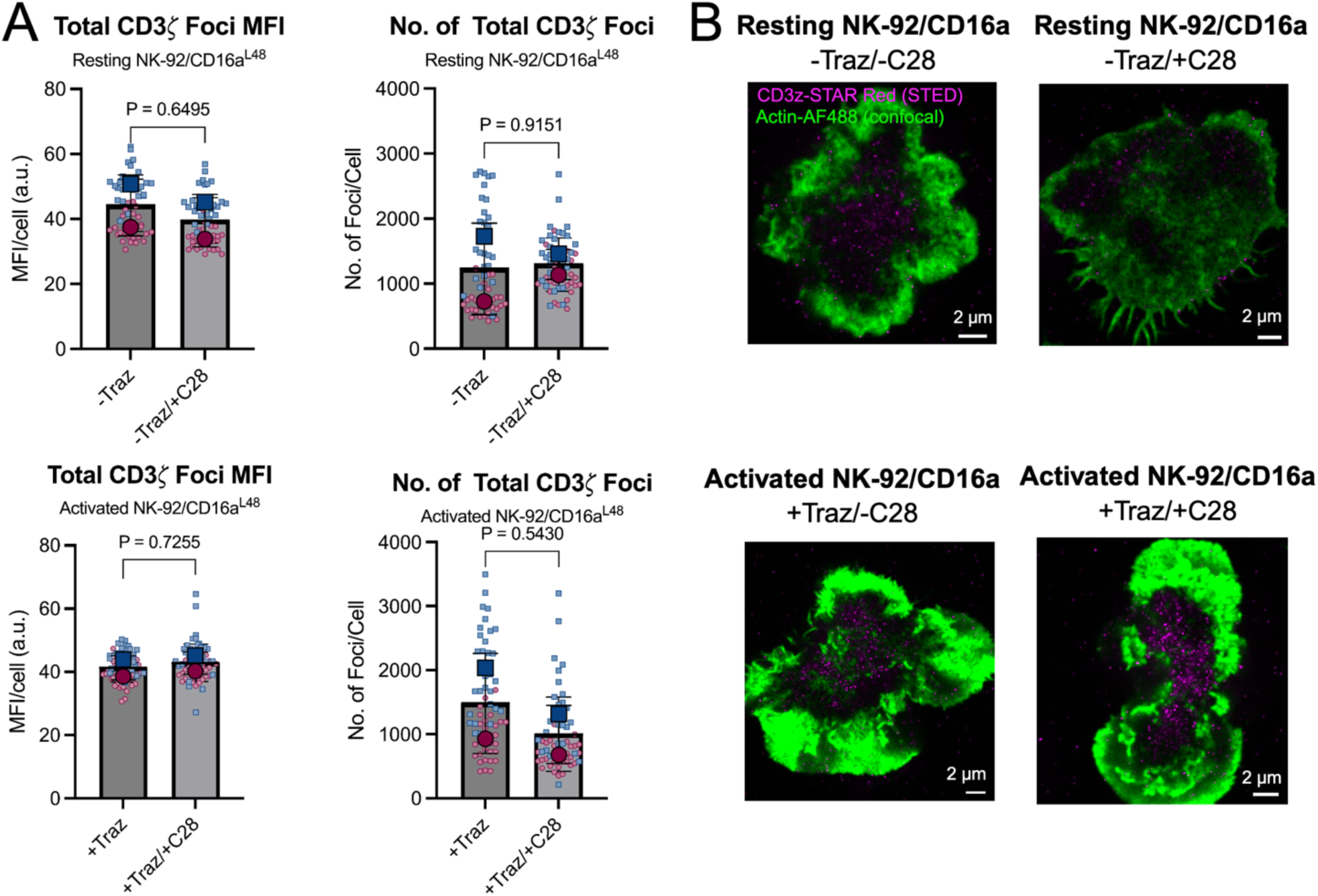
Blocking CD16a D1 does not alter total CD3ζ. A) NK-92/CD16a^L48^ cells were coated on supported lipid bilayers (SLBs) containing ICAM and HER2, with and without C28 (30 µg/mL) and/or Trastuzumab (10 µg/mL) and allowed to form synapses for 10 min before fixing and permeabilizing. Cells were stained for CD3ζ and the number of foci and MFI was quantified. All data are represented using SuperPlots^67^. Two biological replicates were performed (n=60, small symbols) and the mean was calculated (large symbols). Statistical significance was determined using un-paired, lognormal Welch’s t-tests, comparing means of each biological replicate. a.u. = arbitrary unit. B) Representative images of cells from part A, with confocal images of actin (green, phalloidin-488) overlaid upon STED images of CD3ζ (magenta, Star-RED).

### ADCC-enhancing mutations to CD16a D1 do not alter dimer distances

The CD16a variant in D1 at L48H/R is known to enhance ADCC activity^16–18^. Since the L48 residue sits adjacent but close to the C28 epitope (and some L48 contacts are involved), we wondered how the variant L48H affects CD16a dimerization. Since the L48H variant has enhanced ADCC activity over the WT protein, and since we predict the dimer to be the inactive oligomeric state, we hypothesized that CD16a dimers may be wider in H48 versus L48 NK-92/CD16a cell lines due to steric repulsion in the homodimer. We previously utilized MINFLUX nanoscopy to show that CD16a occurs as pairs of proteins on the resting and ADCC activated NK cell surface within the immune synapse^13^. In our previous studies, we utilized a genetically altered NK-92/CD16a cell line with a C-terminal SNAP-tag and localized CD16a using the photoswitching nature of Alexa Fluor 647. However, to more accurately label CD16a, here we used the nanobody C21, which does not interfere with ADCC, IgG binding, or C28 binding. Utilizing a nanobody enables closer labeling to the target protein and enhances specificity. We conjugated a speed-optimized, concatenated DNA points accumulation for imaging in nanoscale topography (PAINT) docking strand sequence onto a C-terminal ectopic cysteine residue using copper-free click chemistry, as previously described^33–35^. We then generated resting cell immune synapses on SLBs as described above using either NK-92/CD16a^L48^ or NK-92/CD16a^H48^ cells without Trastuzumab. Finally, we combined DNA-PAINT with MINFLUX localization to map CD16a with ATTO-647N imager strands (Fig. 6).

**Figure 6.**
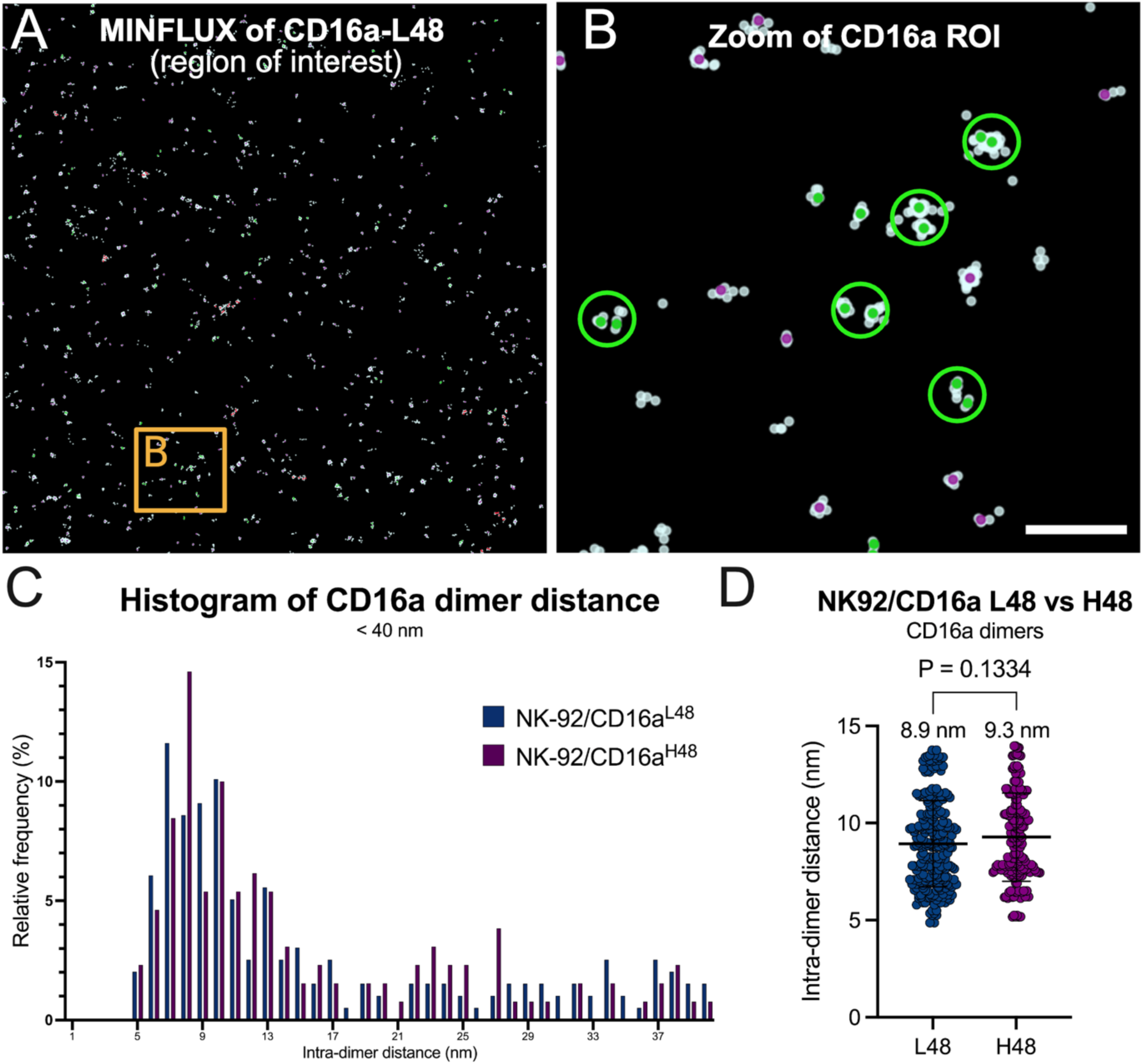
MINFLUX nanoscopy indicates that CD16a D1 isotypes altering ADCC activity do not affect dimer dimensions. A) 2D MINFLUX region of interest (ROI, 3 µm x 3 µm) of CD16a labeled with C21 conjugated to a DNA PAINT docking strand (R1x5) and imaged using a complementary ATTO647N imager strand. Localization clouds (trace) are light blue, and filtered data, indicating the center positions of the individual traces, are colored in purple (single, isolated event), green (dimers, 2 trace centers within 40 nm), and red (clusters of 4 or more traces within 40 nm). B) Zoomed-in region from part A as indicated, showing dimers of CD16a circled in green. C) CD16a intra-dimer distances from both NK-92/CD16^L48^ and NK-92/CD16a^H48^ cells were compiled and plotted in a histogram, indicating a peak in intra-dimer distances centered around 9 nm. D) CD16a intra-dimer distances 15 nm and less were plotted to determine the average distance of CD16a^L48^ and CD16a^H48^ pairs to be 8.9 (+/- 2.2) nm and 9.3 (+/- 2.3) nm, respectively (244 dimers for L48 and 168 dimers for H48). A Welch’s t-test was used to determine statistical significance. For MINFLUX data, n = 5 per condition (each a biological replicate), with at least 5,000 valid localizations collected per experiment. Scale bar = 100 nm.

After filtering (see methods), we detected 1,011 valid trace centers for NK-92/CD16a^L48^ and 714 for NK-92/CD16a^H48^. Of these, 39% and 36% were dimers (2 traces within 40 nm or less), for L48 and H48, respectively. The detection of dimers validated our previous data using an alternative labeling scheme and provides additional evidence that CD16a is likely a dimer in its resting state (Fig. 6A-B). However, it should be noted that data were collected in 2D, therefore it is possible distance measurements are bias due to projection out of the plane of imaging (i.e. dimers whose axis is not parallel to the image plane). However, given the short dimensions and the preferential arrangement of target molecules to be in a common plane, it is likely that the percentage of dimers would be the same when measured in 3D.

Using nearest neighbor analysis, we isolated pairs of CD16a and plotted their distances on a histogram (Fig. 6C). We noted a peak of distances at ∼9 nm, which is much closer than what we previously calculated. This discrepancy can be well explained by the differences in labeling schemes, most importantly the spatially and sterically different epitopes (flexible C-terminus versus D1). The intra-dimer distances of dimers less than 15 nm did not significantly differ between L48 and H48 cells (Fig. 6D), meaning that enhanced ADCC activity of H48 cells is not due to a change in protein oligomerization (at least not one that we can detect at this resolution). These data suggests that CD16a clustering is necessary but not sufficient for ADCC. Further, our data reinforce our previous studies in showing that CD16a is a dimer and that this may be connected to its ability to cluster upon binding to immune complexes within the NK cell immune synapse. However, CD16a D1 could potentiate CD16a dimerization, possibly keeping NK cells from being constitutively active in the presence of such high concentrations of antibodies and immune complexes in the serum.

### Structure-guided mutagenesis to the CD16a D1 C28 epitope enhances ADCC

We next wondered whether eliminating C28 binding to CD16a D1 would enhance ADCC activity. The following hypothesis was guided by our knowledge of the L48H substitution, which also occurs in D1 but in a region adjacent to the C28 binding site and is known to enhance ADCC activity. We hypothesized that C28 may act as a surrogate or outcompete an unknown protein ligand to this epitope that normally must dissociate for signaling to proceed. We reasoned that by knocking out C28 binding, we could eliminate the binding of a potential checkpoint protein that binds in *cis* to the same C28 D1 epitope on the NK cell surface, or that we may disrupt the dimer interface to potentiate more monomers on the cell surface.

Using structure guided mutational analysis, we first designed a set of single mutations that occurred within key residues identified in the C28 epitope. This included D59A, D65A, K114A, and Y56A. We expressed and purified CD16a (V158) with these single point mutations and tested their binding to C28 via BLI. While no single mutation knocked out binding completely, D59A, D65A, and Y56A significantly reduced binding (Fig. 7A). We then combined mutations or tried additional point mutations. Either Y56R alone or combining D59A/W with D65A/W led to a complete knockout of C28 binding (Fig. 7A). We therefore chose to proceed with the Y56R and D59A/D65A mutant variants for analysis of ADCC activity. Notably, the D59A/D65A mutation pair significantly changes the electrostatic potential of the C28 epitope, changing it to a neutral charge (Fig. S5B).

**Figure 7.**
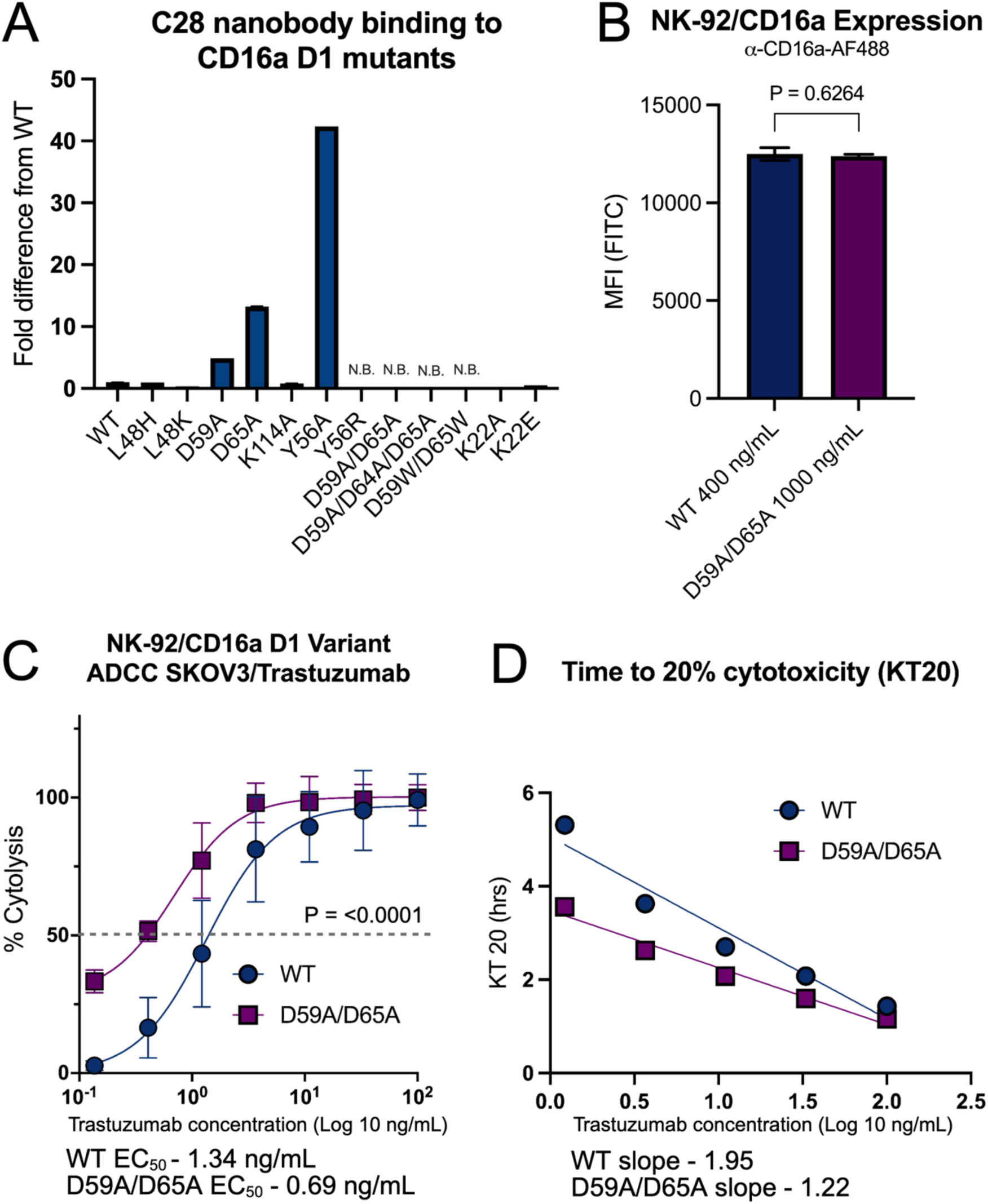
Structure-guide mutagenesis to the C28 epitope on CD16a D1 enhances ADCC activity and killing kinetics. A) CD16a ectodomains with the indicated substitutions, were tested for their binding to C28 using BLI. Higher fold-differences indicate worse binding, while N.B. indicates no binding. B) Lipid nanoparticles (LNPs) containing the indicated CD16a mRNA, titrated at different concentrations, were used to transfect NK-92 cells and CD16a expression was measured using flow cytometry. WT and D59A/D65A constructs had equal surface expression after 24 h at 400 ng/mL and 1000 ng/mL, respectively. C) Cells from part B were used for ADCC assays against SKOV3 cells coated in Trastuzumab. IC_50_ values are indicated and the p-value is the result of a best-fit value comparison of parameters. D) Time to 20% of target cells killed (KT20) was measured across different Trastuzumab concentrations for WT versus D59A/D65A NK-92/CD16a cells. The steeper slope of the D59A/D65A cells indicates faster killing kinetics.

To first assess whether our variants would express in NK cells and to screen them for more detailed downstream analysis, we chose to first transiently transfect NK-92 cells with our variant constructs using mRNA lipid nanoparticle (LNP) delivery. To optimize our LNPs specifically for NK cell transfection, we screened four LNP formulations using EGFP mRNA as a marker of transfection efficiency. Previous optimizations indicate that both substituting traditional cholesterol for beta-sitosterol and modulating the N:P ratio (2.5 – 5) have a significant impact on *in vitro* and *ex vivo* transfection, including in NK cells^36–38^. After characterizing LNP size, polydispersity, and encapsulation efficiency (Fig. S7A), LNPs were added at different concentrations to NK-92 cells and allowed to express for 24h. We found that the formulation of beta-sitosterol LNPs with N:P = 5 (b5-LNPs) gave us the highest transfection efficiency and most robust EGFP expression (Fig. S7B).

We next tested our constructs at 24h and 48h. We found that our mutant constructs showed a lower expression level than the WT construct, with Y56A having the poorest expression (Fig. S7C). Therefore, we chose to move forward with the D59A/D65A variant, with maximum expression peaking around 24h. We first titrated the amount of LNPs we added to target cells to match expression of our variants with that of WT (Fig. 7B). Using these cells, we set up dose-response ADCC assays using SKOV3 cells coated with Trastuzumab, at an E:T ratio of 2:1. We found that the D59A/D65A variant had significantly better EC_50_ value over WT cells (Fig. 6C). Further, the D59A/D65A variant killed target cells much faster and more efficiently, suggesting an enhancement to serially killing (Fig. 6D).

### Structural analysis and molecular dynamics suggest D1 influences CD16a conformation and oligomeric state to regulate activation

A previous study aimed at specifically modulating CD16a activity with affimers found that one such protein, named AfG3, was able to allosterically influence IgG binding^39^. They showed that their affimer stabilized a conformation of CD16a more closely resembling unliganded CD16a. In an Fc-liganded structure, CD16a D1 moves significantly, creating a much wider angle of ∼59° (Fig. 8A). Comparing our structure to unliganded CD16a as well as the AfG3-CD16a structures, we found that C28 bound CD16a also closely resembles the unliganded structure (Fig. 8A). These data suggest that IgG binding may induce a structural change in D1 that allosterically induces an overall conformational change affecting the dimeric, basal CD16a signaling unit. Of note, our structure is the first of CD16a in solution, while all others have been solved in a crystallographic context, which could influence conformation. Also, our structure and others are of isolated CD16a ectodomains and lack the cellular context of the full length signaling complex.

**Figure 8.**
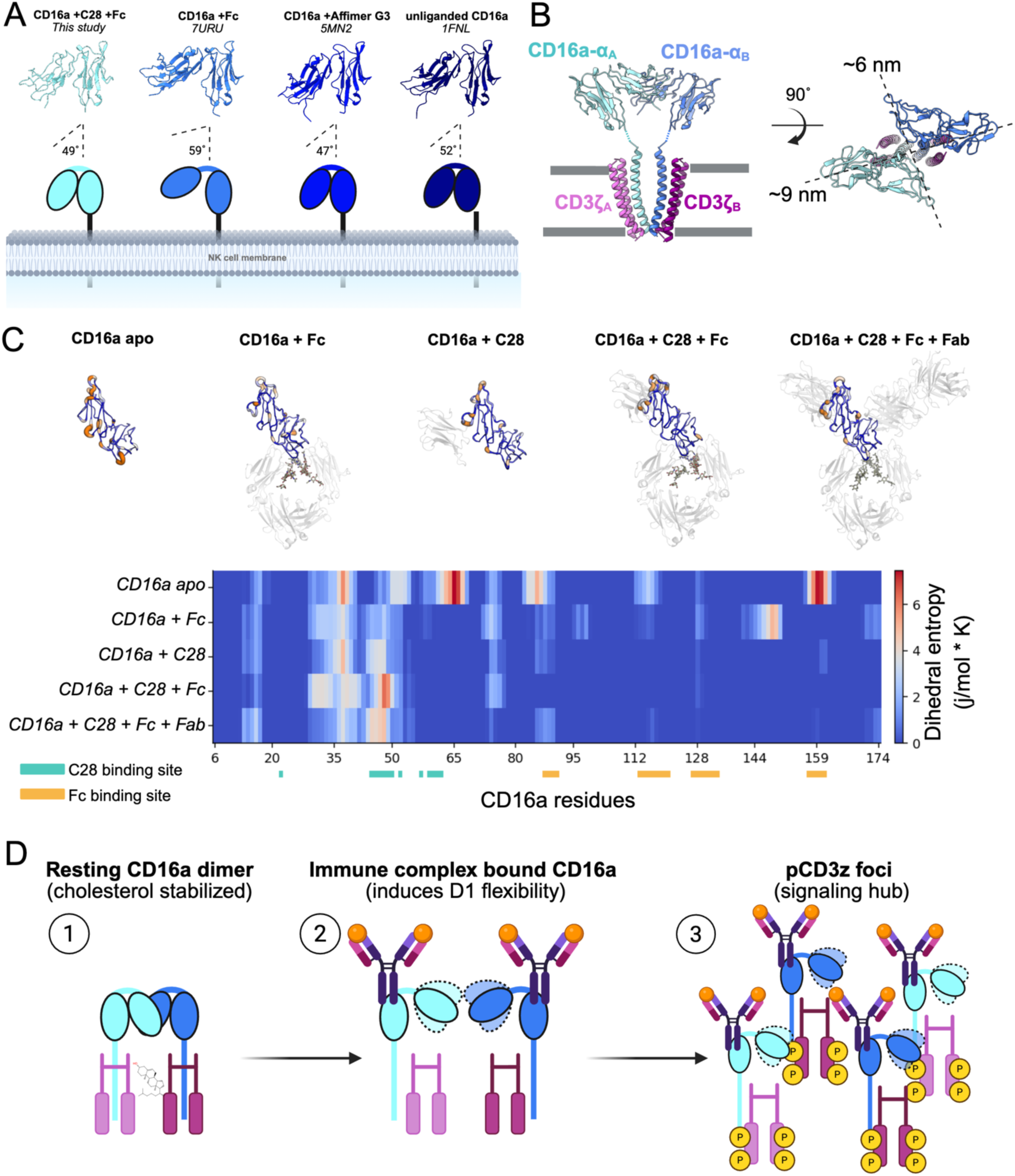
Modeling and molecular dynamics support an overall model of IgG-mediated cytotoxicity in NK cells. A) Comparison of structures of CD16a in different binding states, with a cartoon model below indicating the degree of bend in the ectodomain. The structure of the C28- bound CD16a more closely resembles that of an inactive, affimer-bound structure and that of unliganded CD16a, even with Fc bound. B) The dimer model of the IgE-receptor is shown with our structure of CD16a overlaid on the ectodomains, aligned on the membrane proximal domains. The beta domains were deleted, leaving just the alpha domain and the common gamma chain, which is analogous to the CD3ζ domain (labeled as such). Viewing the dimer from above, such as it is viewed in our MINFLUX data, shows that distances across the dimer range from 6-9 Å, in line with our empirical evidence. C) Molecular dynamics simulations of CD16a bound to different ligands. Larger, warmer regions indicate higher flexibility/dynamics while thinner, cooler regions indicate less motion. Per residue energy landscapes are shown in the chart below for each complex, with the C28 and Fc binding domains indicated. D) Overall working model of IgG-based activation from a resting CD16a dimer: 1) We hypothesize that CD16a, with closed ectodomains, are arranged as dimers and are potentially stabilized by intracellular cholesterol. This oligomeric arrangement prevents kinase access to CD3ζ. 2) Upon immune complex binding, D1 domains become flexible, liberating CD16a monomers bound to immune complexes. 3) Within an immune synapse and with additional activating signals, CD16a forms signaling foci that potentiate phosphorylation of CD3ζ due to kinase access to ITAMs.

Previously, a structure of the Fc epsilon receptor I (FceRI) was solved by cryo-EM and showed that the protein is dimeric, supported by cholesterol within the transmembrane region^20^. The IgE receptor is highly related in function to CD16a and contains many identical structural motifs, including the alpha domain and association with the common gamma chain. It was demonstrated that the dimer was the inactive form, which monomerizes upon IgE binding, leading to activation. Although the extracellular domains of FcεRI were not resolved in the dimer, we modeled IgE monomers onto the dimer structure, matching the alpha TM domains, and then aligned our CD16a structure onto the IgE receptor ectodomains, matching the membrane proximal domains (Fig. 8B). This suggests a potential dimeric motif for CD16a, which we have shown occurs on the surface of NK cells^13^. Several key residues are conserved between the IgE and IgG receptor binding domains, including L48 and V158, and both the IgE and IgG receptors utilize analogous ITAM-containing signaling domains, including the common gamma chain and CD3ζ. In our model, the D1 domains are aligned and interacting near the electronegative patch (Fig. S5), leaving the membrane proximal D2 domains free for IgG binding. However, it is not likely that the electronegative patch serves as the dimer interface. The dimer may be held together by a different patch on the CD16a ectodomain, by an unknown protein in *cis* (as discussed above), via the TM domain protein interfaces and cholesterol, or some combination of the above. Notably, however, the general arrangement of CD16a that we show here agrees with our empirical MINFLUX data in terms of dimensions (Fig. 8B).

We next sought to utilize molecular dynamics (MD) simulations to determine if our hypothesized CD16a D1 motions could be recapitulated. We performed simulations with CD16a by itself, with C28 bound, with solely the Fc bound, with both C28 and the Fc bound and with C28, the Fc and the Fab (P2C-47) present. To quantify changes in dynamics of the CD16a D1 domain, we calculated dihedral entropies (Fig. 8C). The highest degree of flexibility is found for CD16a in its apo state. Binding of the Fc domain stabilizes parts of the Fc epitope and pre-organizes parts of the C28 epitope, however, binding of C28 to CD16a not only results in a more rigid binding interface but also contributes to reduced flexibility close to the Fc epitope (∼residues 130-150). Remarkably, this rigidification extends beyond the direct C28 binding footprint, suggesting that C28 exerts an allosteric effect on distal regions of CD16a. The flexibility that Fc binding introduces is further compensated upon co-binding of C28, and this cooperative rigidification is maintained even in the presence of the Fab fragment. Taken together, these results suggest that C28 is a key stabilizer of CD16a, acting not only locally at its binding interface but also allosterically at the Fc epitope.

Using the data we provide here, and previous work on the analogous IgE receptor, we propose a working model for how CD16a signaling occurs, leading to ADCC (Fig. 8D). We hypothesize that CD16a dimers, like the IgE receptor dimer, represents an inactive state, which holds CD3ζ tightly associated with CD16a TM domains, preventing phosphorylation due to steric hinderance. This state may also be potentially stabilized by the lipid rafts rich in cholesterol. Upon immune complex binding, the D1 conformation shifts and the dimer destabilizes, liberating immune complex-bound monomers to cluster into signaling hubs. The exposed CD3ζ form foci where the ITAMs are then readily phosphorylated by Src and Lck kinases within these signaling hubs. The immune synapse provides co-stimulatory signals, as has been previously demonstrated, that provide polarization that may assist these large shifts in CD16a^5,40–42^. Thus, we propose that since the membrane distal domains of all Ig receptors have a similarly conserved structural motif, this model may be a universal paradigm for Ig-based effector functions through regulating the basal signaling unit oligomeric state.

## Discussion

Here we provide evidence that D1 of the NK cell activating receptor CD16a functions as a checkpoint on ADCC-based activation through allosteric influence of oligomeric shifts and ITAM-based signaling, independent of IgG binding. Using a pair of nanobodies with non-overlapping D1 epitopes, we show that occupancy of a discrete electronegative patch in D1 (defined by the C28 epitope) is sufficient to block ADCC without disrupting direct cytotoxicity or IgG-Fc binding to the membrane-proximal D2. A cryo-EM structure of CD16a in complex with IgG Fc and both C21 and C28 nanobodies, together with biolayer interferometry, STED-based phospho-signaling analysis, MINFLUX nanoscopy, molecular dynamics simulations, and structure-guided NK cell engineering, collectively support a model in which the dimeric CD16a signaling complex is held in a tonically inactive configuration, and in which immune-complex binding liberates this constraint to enable downstream CD3ζ phosphorylation and cytolysis.

The functional distinction between nanobodies C21 and C28 is striking and was not anticipated from their shared D1 binding location. Both nanobodies bind exclusively to D1 and do not overlap with the IgG-Fc binding site in D2, yet only C28 inhibits ADCC. The C28 epitope is centered on top of CD16a D1, overlapping with an electronegative surface that includes D59 and D65, and its CDR3 makes an additional bridging contact with K326 of IgG Fc (Fig. 4F-G). This single inter-chain hydrogen bond and salt bridge in the ternary complex may not be sufficient to account for steric competition with Fc, yet biolayer interferometry demonstrates that C28 actively displaces core-fucosylated IgG and prevents its re-association (Fig 2B-C). Critically, the block is allosteric rather than purely steric: afucosylated antibodies, which engage CD16a D2 with markedly higher affinity, can rebind in the presence of C28 and partially recover biosensor signal (Fig. 2F-G), yet ADCC is still attenuated under these conditions (Fig. S2). This dissociation between binding competence and effector function activation points to a mechanism in which D1 does not simply serve as a steric gatekeeper for Fc access but instead transduces conformational information across the ectodomain to regulate the intracellular signaling-competent state of the receptor complex. Consistent with this interpretation, comparison of our cryo-EM structure to previously published crystallographic and affimer-bound CD16a models shows that C28-bound CD16a adopts a closed D1 conformation indistinguishable from unliganded receptor, even in the presence of Fc, whereas Fc alone induces a wider inter-domain angle (Fig. 8A). Molecular dynamics simulations further support the notion that D1 acquires increased conformational flexibility upon Fc engagement, a dynamic mode that is suppressed by C28 binding. Together, these observations suggest that productive ADCC activation requires a Fc-induced change in D1 dynamics that C28 prevents, rather than simply requiring Fc occupancy of D2.

The identification of this allosteric regulatory site is consistent with, and extends, prior work on CD16a function. The monoclonal antibody B73.1, which targets D1 and blocks ADCC without displacing IgG, has been used for decades as a functional tool but its epitope and mechanism remained unresolved^15^. The C28 epitope defined here likely overlaps substantially with or lies adjacent to the B73.1 binding site, providing a long-awaited structural explanation for the paradoxical ability of D1-targeting reagents to block ADCC independently of IgG binding. Separately, the naturally occurring L48H/R polymorphism in D1 is known to abrogate CD2 binding and enhance ADCC by reducing immune synapse size and promoting serial killing^22^. Residue L48 contributes hydrophobic contacts to the C28 epitope and our data show that the L48H substitution modestly reduces C28 sensitivity, though it does not abolish it. Importantly, DNA-PAINT MINFLUX nanoscopy with C21 (which does not perturb ADCC) demonstrates that CD16a forms dimers with ∼9 nm intra-dimer distances on the NK cell surface and that this distance is statistically indistinguishable between L48 and H48 variants (Fig. 6). The ADCC-enhancing effect of the L48H/R substitution therefore does not operate by altering gross receptor dimerization geometry, pointing instead to subtle allosteric changes within D1 that destabilize inhibitory inter-domain contacts. Our MINFLUX measurements also refine our previous estimates of CD16a dimer spacing, which were inflated by the positioning of the SNAP-tag at the flexible C-terminus; the ∼9 nm distances measured here using the compact C21 nanobody as a fiducial are fully consistent with a head-to-head dimer model in which opposing D1 domains contact one another across the synaptic interface (Fig. 8B).

The signaling consequences of C28-mediated D1 engagement were most directly revealed by imaging analysis of pCD3ζ foci in immune synapses formed on supported lipid bilayers and glass. C28 significantly lowers pCD3ζ accumulation in response to Trastuzumab and Rituximab, and notably also reduced basal pCD3ζ in resting NK cells in the absence of antibody, suggesting that D1 exerts tonic inhibitory control over the associated signaling machinery (Fig. 4). This suppression is consistent with a model in which the dimeric receptor organization at rest sterically restricts kinase access to CD3ζ ITAMs. The proximal position of CD3ζ relative to the transmembrane domain of CD16a^28,30,32^ and the documented association of the signaling complex with cholesterol-rich membrane domains^42^ support the idea that receptor dimerization immobilizes CD3ζ within a lipid environment unfavorable for kinase-mediated phosphorylation. The fact that C28 inhibits ADCC even in YTS cells, which lack CD2, confirms that neither CD2 co-engagement nor natural cytotoxicity pathways are responsible for this effect: the D1 epitope acts cell-autonomously within the CD16a signaling unit. Notably, CD3ζ organization shifts in the immune synapse during activation, likely through antibody-induced clustering changes and polarization, but CD16a signaling is still attenuated by C28 binding to D1 despite no change in total CD3ζ (Fig. 5), suggesting that blocking CD16a D1 only prevents kinase access but not the overall CD16a-associated CD3ζ re-organization accompanying ADCC.

Parallels with the structurally related IgE receptor FcεRI lend broader mechanistic weight to our model. A recent cryo-EM structure of the intact FcεRI complex revealed that the receptor is organized as a cholesterol-stabilized dimer in its inactive form, with the α subunits likely held in a configuration that prevents IgE engagement, and that monomerization is required for activation^20^. Although the extracellular domains of FcεRI were not resolved in the dimeric structure, alignment of our CD16a ectodomain onto the FcεRI α chain places the two D1 domains face-to-face near an electronegative patch defined by the C28 epitope (Fig. 8B). Key residues are shared between the IgG and IgE receptor families, and both utilize analogous ITAM-bearing signaling subunits. This structural and functional convergence suggests that membrane-distal Ig-like domains function to autoinhibit receptor signaling through homo-oligomerization. This may be a general organizational principle among activating Fc receptors rather than a peculiarity of CD16a. If so, the regulatory epitope described here could represent a conserved target class for modulating effector cell activation across allergic, autoimmune, and oncologic contexts.

Guided by our structural data, we demonstrated that targeted mutagenesis of the C28 epitope abolishes C28 binding while preserving CD16a surface expression. The electronegative patch that overlaps with the C28 epitope (Fig. S5) echoes other signaling protein charged patches that are thought to also play a role in tonic signaling for effector functions, including by influencing protein-protein interactions and clustering^43^. When delivered as mRNA in LNPs optimized for NK-92 cell transfection, CD16a D59A/D65A conferred significantly improved ADCC potency (lower EC₅₀) and accelerated cytolysis kinetics relative to the wild-type receptor (Fig. 7C-D). Our structural and modeling data suggest that this D1 mutation may favor a more monomeric state of CD16a (Fig. 8C, Fig S5), accelerating ADCC activation kinetics and thus enhancing antibody potency. This phenotype is comparable to those reported for the L48H/R variant but is likely mechanistically distinct. Future development of the C28 epitope as an NK cell engineering strategy will reveal if there are better optimized mutations that can enhance ADCC even further, while preserving natural cytotoxicity (in contrast to the L48H/R variant). Our LNP-mediated transient expression strategy also validates a platform for rapidly screening receptor variants in primary-like NK cell backgrounds prior to stable integration, which may accelerate the preclinical development of next-generation NK cell products.

Several important questions remain. The identity of a potential putative endogenous ligand (or ligands) that engage the electronegative C28 epitope in *cis* on the NK cell surface, and the location of the homo-dimer interface on CD16a, is unknown. CD2 is an attractive candidate given its known interaction with CD16a D1 near L48, but our data showing that C28 blocks ADCC in CD2-negative YTS cells and that addition of soluble CD2 ectodomain does not rescue activity argue against CD2 as the sole relevant contact (Fig. S1B-C). An unidentified inhibitory co-receptor, a serum protein competing for the D1 site under physiological antibody concentrations, or a symmetric D1-D1 contact within the dimer itself are all plausible alternatives that warrant investigation. Additionally, while our biophysical and structural studies used the soluble monomeric CD16a ectodomain, the cellular receptor operates in the context of a dimeric signaling complex, a lipid bilayer, and a crowded immune synapse. How CD3ζ and CD16a are linked within the NK cell surface, and what happens post-immune complex engagement, should also be investigated using multiplexed single molecule imaging methods. Whether the allosteric linkage between D1 and the IgG binding site is fully recapitulated in the full-length receptor, and how membrane curvature and lipid composition modulate the transition between active and inactive dimer states, remain open questions best addressed through cell-based structural approaches. Finally, the potential of D1-targeted interventions to modulate ADCC in primary human NK cells across diverse genetic backgrounds, including the V158/F158 polymorphism that influences IgG affinity and clinical response to mAb therapy, deserves systematic evaluation.

In summary, our data support a model where D1 of CD16a acts as an allosteric regulatory module that couples receptor dimerization to the control of ADCC. By defining the structural basis of this regulation, demonstrating its functional consequence at the level of pCD3ζ signaling, and showing that targeted disruption of the key epitope enhances NK cell killing, we provide both a mechanistic framework and a practical engineering blueprint for improving antibody-based cancer immunotherapies. The parallels with FcεRI suggest that this paradigm may extend broadly across the Fc receptor family, with implications for the rational design of enhanced cellular therapeutics and for understanding the dysregulation of Fc receptor signaling in autoimmune and allergic disease.

## Materials and Methods

### Cell lines and cell culture

SKOV-3 cells (HTB-77), Raji cells (CCl-86) and NK-92 cells (CRL-2407) were purchased from ATCC (Manassas, Virginia, United States of America). SKOV-3 cells were grown and passaged in DMEM (Gibco) media supplemented with 10% (v/v) FBS (Gibco) and 1X Antibiotic-Antimycotic (Anti-Anti, Thermo Fisher). Raji cells were maintained in RPMI-1640 medium with GlutaMAX (Gibco) supplemented with 10% (v/v) FBS and 1X Anti-Anti. NK-92 cells were grown and passaged in MyleoCult (STEMCELL Technologies) media supplemented with 10% FBS, 1X Anti-Anti, and 200 IU/mL IL-2 (Peprotech). IL-2 was added fresh from thawed aliquots each passage, kept at -80°C. Cells were maintained for no more than 4-6 weeks and regularly tested for mycoplasma contamination.

NK-92/CD16a cells came from two sources. For initial ADCC assays and C28 blocking assays (Fig. 1A-B), NK-92/CD16a cells that has been CRISPR-engineered by activating the endogenous CD16a locus were used^21^. For subsequent assays, NK-92/CD16a cells (V158 and either L48 or H48 isotypes) that had been generated by retroviral^44,45^ transfection was used. YTS All cells were maintained identically to parental NK-92 cells.

### Isolation of primary NK cells

For C28 blocking assays, PBMCs were first isolated from whole blood using a SepMate PBMC Isolation spin column, according to the instructions (STEMCELL Technologies). NK cells were then isolated using an EasySep Human NK Cell Isolation Kit (STEMCELL Technologies) according to the instructions and used immediately.

For glass activation assays, primary NK cells were isolated from peripheral blood by negative selection using RosetteSep human NK enrichment cocktail (Stemcell Technologies, 15065). The blood was then layered over a Ficoll–Paque (Thermo Fisher Scientific, 45-001-750) density gradient and centrifuged at 1200 *g* for 20 min without applying a brake. NK cells were harvested from the interface of the gradient and transferred into PBS at room temperature (RT), followed by a 10 min centrifugation at 300 *g* with a low brake. The resulting cell pellet was resuspended in 20 ml of warm R10 and centrifuged again at 1200 *g* for 10 min, also with a low brake. After discarding the supernatant, the cells were resuspended in R10 at a concentration of 2.5×10^6^ cells/ml. NK cells were then cultured for 2 days in low dose IL-15 (5 ng/mL).

### Generation of recombinant proteins

To produce recombinant CD2 ectodomains (eCD2), a mammalian expression for CD2 (SinoBiological) containing a stop codon after residue 206 (corresponding to the topological domain, UniProt P06729) was used to transfect Expi293F (Gibco) cells according to the manufacturer’s protocol. Affinity purification was performed using a 5 mL HisTrap HP column (Cytiva) and was further purified by size-exclusion chromatography (Akta Pure, Cytiva) using and S200I column (Cytiva).

Unless otherwise indicated, IgGs for ADCC assays, including Trastuzumab, Cetuximab, Pertuzumab, and Rituximab were ordered from SelleckChem. Trastuzumab, Cetuximab, and Pertuzumab used for fucose/fucose-free assays, as well as PGT-121, were produced by recombinant protein expression in mammalian cells. Heavy chain (HC) and light chain (LC) sequences for these antibodies were synthesized and cloned into pCDNA3.4 (Genscript). Then, plasmid DNA at a 2:1 HC:LC ratio was used to transfect ExpiCHO (Gibco) cells according to the manufacturer’s protocol. For fucose-free antibodies, 2-deoxy-2-fluoro-fucose was added at 250 µM final concentration directly to cultures at the same time as transfection. This has been reported to reduce fucosylation by >90%^46^. Affinity purification was performed using a 5 mL PrismA column (Cytiva) followed by size exclusion chromatography (SEC) using an S200I column.

An expression plasmid encoding the V158 variant of the CD16a ectodomain (containing a C-terminal 6-His tag and glycan deletions N38Q, N74Q, and N169Q) was obtained from Pamela Bjorkman^26^. CD16a was recombinantly expressed in the presence of kifunensine (2.5 µM) and purified from HEK 293F cells as described above (750 µg of DNA per liter of cells). An expression plasmid encoding the human Fc domain (V158) was obtained from Adam Barb^46^ and used to express and purify Fc as described for IgGs above, but in HEK 293F cells (750 µg DNA per liter of cells). Nanobodies C21 and C28 used for structural studies were also expressed and purified by mammalian expression in HEK293F cells (750 µg DNA per liter of cells). Sequences were synthesized (Genscript) and cloned into the expression vector AbVec2.0-IGHG1 plasmid (Addgene #80795), utilizing a C-terminal double streptavidin tag. Proteins were affinity purified using a 5 mL StrepTrap XT column (Cytiva) followed by SEC on an S75I column (Cytiva). The HC and LC sequences for the P2C-47 Fab were synthesized and cloned into AbVec2.0-IGHG1 and AbVec1.1-IGKC (Addgene #80796), respectively (Genscript). Fab was expressed in Expi293F cells as described above and affinity purified using a CH1-XL column (ThermoFisher), followed by SEC using an S75I.

### Flow cytometry

All flow cytometry was performed using a CytoFlex Flow Cytometer (Beckman Coulter). For evaluating GFP expression, 200 µL of cells were added to a round bottom 96-well plate and centrifuged at 300 x g for 5 min, washed once in 1X PBS + 1% (v/v) FBS, and then resuspended in 200 µL PBS/FBS. The 488-nm FITC laser was used, and negative gating was set with parental NK-92 cells. For CD16a expression, a similar approach was utilized. After washing, cells were stained with 1 µL of anti-human CD16a-AF488 (3G8, Biolegend) for 1h on ice, before being washed a second time. For gating, the 488-nm FITC laser was again used and the positive gate was set using NK-92/CD16a^L48^ cells mixed 1:1 with NK-92 parental cells.

### Competition assays using BLI

For BLI competition studies, a WT version of CD16a with an N-terminal 6-His tag and a C-terminal Avi-tag was utilized (synthesized, cloned, expressed and purified by Genscript). Nanobodies C21 and C28 utilized for competition assays were synthesized, cloned, expressed and purified by Genscript. Nanobodies were expressed in *E. coli* and contained a C-terminal 8-His tag and Avi-tag. Antibodies used for competition assays were ordered from SelleckChem, while PGT-121 was made in-house (see above). All BLI assays were performed using an Octet RED96e system (Sartorius). Strepavidin (SA) biosensors (Sartorius) were first activated in 1X Kinetics Buffer (KB, 1X PBS, 0.1% v/v Tween-20, 1% BSA) for 30 s. Biotinylated CD16a was then loaded onto tips at 50 µg/mL in 1X KB until 2 nm binding was achieved. Tips were then washed for 60s in 1X KB followed by association in either IgG or nanobody (5 µM) in 1X KB for 60s. Tips were then dipped into solutions containing an equal mixture of IgG and nanobody to maintain the concentration of the previous analyte while introducing a second competitor. Raw sensogram data were plotted using Prism 11.

### Site-directed mutagenesis binding assays using BLI

For binding kinetics studies, nanobodies used for competition assays (see above) were first biotinylated using a kit (Avidity LLC). All CD16a WT and mutant constructs (on V158 background) were synthesized, cloned, expressed and purified using a 6-His tag by Genscript in mammalian cells. SA biosensors were first activated in 1X KB for 30 s. Biotinylated nanobodies were then loaded onto tips at 50 µg/mL in 1X KB until 2 nm binding was achieved. Tips were then washed for 60s in 1X KB followed by association and dissociation steps of 60 s each in serially diluted CD16a in 1X KB. Starting concentrations of CD16a were determined by optimization around the calculated WT dissociation constant (K_D_). Fold-differences in binding were determined by comparing to the WT CD16a construct K_D_.

### ADCC assays

ADCC assays were completed using the Agilent xCELLigence RTCA system, which uses impedance to measure cell death in real time. For immunotherapy assays, E-Plate 96 PET plates were utilized. A bank measurement was first conducted with 50 µL media (complete RPMI) before 7000 SKOV3 cells per well were added. Plates were allowed to equilibrate at room temperature (RT) for 30 min. Next, target cells were allowed to attach overnight at 37°C/5% (v/v) CO_2_. The following day, target cells were coated in serially diluted antibody in a 50 µL volume of complete media. When nanobodies were added, a constant concentration of antibody was first added in 25 µL media, followed by 25 µL of serially diluted nanobodies. Finally, effector NK cells were then added at the appropriate E:T ratio in a volume of 50 µL media. After equilibrating for 30 min at RT, plates were added to the RTCA instrument, and the cell index was measured every 15 min for the duration of the experiment. To normalize ADCC assays, a control of just effector cells and target cells was included. For C28 blocking assays, a control of just effector and target cell with antibody alone was used and values were reported as either percent cytotoxicity or normalized cell index. All analyses were completed using the Agilent RTCA software. Nanobodies C21 and C28 utilized for ADCC assays were the same as those used for competition and binding assays.

### Cryo-EM sample prep

To form the complex for cryo-EM, 300 µg of CD16 (triple glycan mutant, treated with kifunensine) was mixed with 5M excess of Fc (afucosylated), C21, C28, and P2C-47 Fab. All proteins had been purified on SEC prior into 1X Phosphate Buffered Saline (PBS), pH 7.4. The complex was allowed to equilibrate overnight at 4°C and then purified by SEC using an S200I column (in 1X PBS, pH 7.4). The complex was then concentrated in a 0.5 µL, 100kDa MWCO Amicon Ultra Centrifugal Filter (Millipore) to 1 mg/mL. Directly before vitrification, 6 µL of sample was mixed with 1 µL of n-Octyl-beta-D-glucopyranoside (OBG) that was at 7x the critical micelle concentration of 25 mM. Two grids 1.2/1.3 UltraAuFoil grids were then vitrified in nitrogen cooled ethane using a Vitrobot Mark IV (Thermo) at 4°C, 100% humidity, with a 4 s blot time and 10 s wait time (blot force 0).

### Cryo-EM data collection and processing

Data was collected on Thermo Fisher Glacios, operating at 200 keV mounted with Thermo Fisher Falcon 4 direct electron detector, using the Thermo Fisher EPU 2 software at a magnification of 190,000x. Micrographs were collected at a total cumulative dose of 60.00 e-/Å2 (7.259 pe/p/s) with a pixel size of 0.725 Å, with a 4.34 second exposure time. CryoSPARC Live Patch Motion Correction was used for alignment and dose weighting of movies. CTF estimations, particle picking, particle extraction, iterative rounds of 2D classification, ab-initio reconstruction, heterogeneous refinement, homogeneous refinement, 3D Variability, local refinement, and non-uniform refinement were performed on CryoSPARC. The processing workflows for each dataset is shown in Supplementary Figures S2.

### Model building and refinement

Initial model building was performed by utilizing existing starting models for afucosylated Fc (PDB XXXX), CD16a (PDB XXXX) and P2C-47 (PDB XXXX). For nanobodies, starting models were generated using AlphaFold 3. Models were manually docked into EM density using UCSF Chimera^47^ and ChimeraX^48^. An initial round of refinement was performed using Phenix real space refine, followed by iterative rounds of manual adjustments in Coot followed by refinement in Phenix to reach the final model. Contact analysis was performed using PLIP^49^, and figures were created in UCSF Chimera and ChimeraX.

### Lipid nanoparticle preparation and transfection

In vitro transcription of mRNA was performed with in-house T7 using a polyadenylated amplicon template and included 100% uridine substitution by N1-methylpseudouridine (TriLink BioTechnologies) and cotranscriptional capping by CleanCap® Reagent M6 (TriLink BioTechnologies) according to manufacturer’s directions. Following amplicon template digestion (37°C, 30 min, TURBO DNase, Invitrogen), transcripts were purified using carboxylic acid Dynabeads (Invitrogen). Transcript integrity was verified by TapeStation analysis (Agilent) according to manufacturer’s instructions. Capping was confirmed by DNAzyme-facilitated cleavage within the constant 5’ UTR region, between the 24^th^ and 25^th^ templated ribonucleotide (method adapted from Vlatkovic et al^50^; DNAzyme designed using DNAzymeBuilder, https://iimcb.genesilico.pl/DNAzymeBuilder). Resulting RNA fragments were size-separated by 21% Urea-PAGE electrophoresis and capping was evaluated by fragment migration compared to capped and uncapped controls.

LNPs were produced using SM-102 (Cayman Chemical), 1,2-distearoyl-sn-glycero-3-phosphocholine (DSPC; Avanti Polar Lipids), 1,2-Dimyristoyl-rac-glycero-3-methoxy-Polyethylene Glycol(2000) (DMG-PEG2000; Cayman Chemical), and either cholesterol (Cayman Chemical) or beta-sitosterol (MedChemExpress) at a molar ratio of 50:10:1.5:38.5 respectively, with a final total lipid concentration of 6.2 mg/mL. mRNA in 100 mM NaOAc, pH 5.2 was mixed with lipids in ethanol at a 3:1 ratio, 12 mL/min total flow rate, using a NanoAssemblr Ignite (Precision Nanosystems). mRNA concentration was determined by NanoDrop (ThermoScientific) and adjusted for desired nucleic acid (N) to phospholipid (P) ratio. Resulting LNPs were buffer-exchanged 2500-fold into sterile DPBS by sequential spin-filtration at 2,000 x g in a 10 kDa cutoff Amicon spin filter (Millipore). Encapsulation efficiency and cargo concentration were determined by standard Quant-iT™ RiboGreen assay (ThermoFisher Scientific). Concentrations were normalized and 1 volume of 10% sucrose in DPBS was added (final concentration 5% sucrose). Z-average diameter and polydispersity index were evaluated by dynamic light scattering (Unchained Labs). LNPs were aliquoted and frozen at a rate of ∼1°C/min and stored at -80°C until use.

To transfect NK-92 parental cells for LNPs, cells were washed and resuspended in 1.5 mL serum-free RPMI. LNPs, prepared at 20 ng/mL, were thawed rapidly in a water bath at 37°C just until almost completed thawed, then briefly vortexed before being added to cells at the desired concentration. Cell/LNP mixtures were divided among three wells of a 12-well plate and allowed to incubate at 37°C/5% CO_2_ for 2h before 500 µL of complete Myleocult media + IL-2 (200 IU/mL) was added. Cells were allowed to incubate 24-48 prior to utilizing in flow cytometry or ADCC assays.

### Liposome preparation for SLBs

Liposomes for SLBs were prepared as previously described^13^. Briefly, liposomes were formed by dissolving 1-palmitoyl-2-oleoyl-glycero-3-phosphocholine (POPC) and 1,2-dioleoyl-sn-glycero-3-[(N-(5-amino-1-carboxypentyl)iminodiacetic acid) succinyl] (nickel salt) (DGS-NTA(Ni)) (Avanti) in chloroform at a 96:4 molar ratio. Lipids were dried under vacuum overnight, hydrated, and then sonicated for 30 s. Liposomes were then extruded sequentially through 0.8-μm–0.1-μm filters using a Mini Extruder (Avanti) at RT.

### Light microscopy sample preparation

To prepare NK cell synapses for STED microscopy imaging, slides were prepared as described previously^13^. Briefly, SLBs were prepared on cover glasses (Deckgläser coverslips, #1.5) that were thoroughly cleaned. Coverslips were then oxygen plasma (Solarius) treated before being attached to a 6-channel sticky-slide (Ibidi VI 0.4). Liposomes were deposited into each channel, incubated for 20 min at room temperature, and thoroughly washed with PBS. SLBs were incubated with 100 µM NiCl2 containing 1% (w/v) BSA for 20 min at room temperature, washed, and then incubated with His-tagged HER2 (10 µg/mL; Acro Biosystems) and His-tagged ICAM-1 (1 µg/mL; Acro Biosystems) for 60 min at 37 °C, and then washed with PBS. SLBs were then incubated either alone or with Trastuzumab (Selleckchem) at 10 µg/mL for at least 30 minutes at room temperature, washed with PBS, then chambers were finally left to sit in PBS or PBS containing 30µg/mL C28 nanobody during cell preparation. Next, 6 × 10^6^ NK cells, either alone or incubated with 30 µg/mL C28 nanobody for at least 30 min, were applied and incubated at 37 °C in 5% CO_2_ for 10 min. Samples were then washed, fixed with 4% (w/v) PFA, permeabilized, and post-fixed/quenched with 4% PFA. Actin was labeled with phalloidin-AF488 (Invitrogen), diluted 1:400 from a 66 µM stock in DMSO, and pCD3ζ or total CD3ζ was simultaneously probed with a human α-pCD3ζ antibody (BD Biosciences) or with a human α-CD3ζ antibody (Biolegend, clone 6B10.2) at 5 µg/mL at 4 °C overnight. The next day, cells were washed with 1X PBS and counterstained with anti-mouse STAR RED (Abberior) at 5 µg/mL for 1h at 4°C, followed by a second wash prior to imaging.

For confocal imaging of immune synapses on glass, primary NK cells were incubated for 20 minutes at 37°C with 5% CO2 on glass chamber slides pre-coated with 1.25 μg/ml Fc-ICAM-1 (Enzo Life Science; ALX-203-004-C050) with or without 5 μg/ml rituximab (Selleckchem; A2009). Cells were fixed at RT for 15 min with freshly prepared 4% paraformaldehyde (Thermo Fisher Scientific; 50-980-487) in 1X PBS and permeabilized with 0.25% Triton X-100 (Thermo Fisher Scientific; AAA16046AE) at RT for 5 min. Next, 10% goat serum (Sigma-Aldrich; NS02L) was added to block non-specific binding followed by 1mg/ml of sodium borohydride (Sigma-Aldrich; 71320) to quench auto-fluorescence. Phalloidin Alexa Fluor 568 (Thermo Fisher Scientific; A12380), phospho-CD247 (pY142) were used for staining. Coverslips were rinsed 3 times with 200 μl of 1X PBS between each step. Images were captured using a 100× 1.46 NA objective on a Zeiss AxioObserver Z1 microscope stand fitted with a Yokogawa W1 spinning disk.

### Light microscopy data collection and processing

STED images of NK cells were collected on an Abberior Facility Line STED instrument (Göttingen, Germany), using an Olympus 60x UPLanXApo 1.42NA objective. Actin was imaged using phalloidin with 20% 488nm laser with an emission window of 498nm-608nm. pCD3ζ was imaged using STAR RED secondary antibody with 10% 640nm laser, and 11.2% 775nm depletion laser power with 6 accumulations. The emission window for STAR RED was 650nm-755nm. Pixel size was 30nm with 5µsec dwell time. Plane of focus was individually set at the synapse surface for each cell with the 488nm laser using the ‘Live’ imaging mode. 30 cells from each of the 4 treatment conditions were imaged. Raw images were exported and processed using Imaris software (Version 10.2). Confocal actin images were used to create a mask of each cell outline using the ‘surfaces’ module. pCD3ζ puncti inside of the mask area were recognized and analyzed for mean fluorescence intensity (MFI), number, and area (as number of voxels). This was done using two methods: manually thresholding the parameters in the surfaces module or using a machine learning algorithm. Statistics for MFI, number, and area of pCD3ζ puncti for each cell were exported as .csv files then compiled, and averages obtained using a custom python script. Statistical analysis was performed in Prism 11 software. Exemplary images from raw images were created using Fiji software (Version 2.14.0).

### DNA-PAINT nanobody probe preparation

R1 x 5 DNA oligonucleotide docking strands with a 5’ azide and corresponding imager strands containing a 3’ ATTO647N fluorophore were ordered from IDT. Nanobody C21 containing an N-terminal ectopic cysteine was expressed and purified as described above. To conjugate docking strands to C21, click chemistry was performed. Nanobodies were suspended in 1× PBS containing 5 mM TCEP (Bond Breaker TCEP, pH 7.0) and incubated with a 10-fold molar excess of maleimide-DBCO for 5 hours at room temperature, followed by overnight incubation at 4°C. Excess maleimide-DBCO was removed by three sequential passes through Zeba Dye and Biotin Removal Spin Columns (ThermoFisher) that were first equilibrated in 1× PBS. Protein concentration was re-measured and then azide-functionalized oligonucleotides were added at a 10-fold molar excess and reacted under the same conditions (5 hours at room temperature, overnight at 4°C). Conjugated nanobody-oligonucleotide complexes were purified by size-exclusion chromatography on a Superdex 75 Increase 10/300 column; the conjugated species was identified by A260/A280 ratio. Purified conjugates were flash-frozen and stored in aliquots at −80°C.

### Cell Labeling and Sample Preparation for DNA-PAINT MINFLUX Nanoscopy

Nanobody C21 bearing an azide-linked DNA-PAINT docking strand (R1 x 5) was diluted to 50 nM in labeling buffer consisting of PBS supplemented with 1% (w/v) BSA and 1 mM DTT. Cells were incubated in 100-200 µL of labeling solution for 50 minutes at room temperature, or overnight at 4°C. Labeling solution was discarded after use. Cells were then washed twice with PBS for 5 minutes per wash, followed by a final wash for 30 minutes.

Cells were next post-fixed with 4% (w/v) paraformaldehyde (PFA) for 15 minutes at room temperature and quenched with 50 mM NH₄Cl for 5 minutes. Samples were washed four times with PBS (5 minutes per wash). For samples not imaged within a few days, PBS was supplemented with 0.02% (w/v) sodium azide for storage at 4°C.

On the day of imaging, an imaging buffer was freshly prepared at containing 10 mM MgCl₂, 5 mM Tris-HCl (pH 8.0), 1 mM EDTA, and 0.05% (v/v) Tween-20. PBS was exchanged for imaging buffer, leaving sufficient volume to keep the imaging chamber covered. Gold nanoparticles in the 2-3 nm size range (AUROlite^TM^ Au/TiO_2_, Strem), used as fiducial markers for drift correction, were resuspended and 100 µL was applied to each chamber for 10 minutes at room temperature. Unbound gold was removed by washing several times with imaging buffer. Immediately prior to imaging, imaging buffer was replaced with ATTO647N-conjugated imager strand diluted to 200 pM in imaging buffer.

### MINFLUX data collection

All MINFLUX data were acquired on a commercial MINFLUX microscope using the Imspector software with MINFLUX drivers (Abberior Instruments). The sample position in relation to the microscope was actively stabilized on the backscattering of a 975-nm-widefield illumination by the gold beads (msd < 3 nm for all 3 axes)^51^. Immune synapses at the plane of the bilayer were identified visually using the 488 nm laser in confocal mode. Next, regions of interest (ROIs) of ∼3 µm x 3µm were selected. In confocal mode, the z-height of the sample was adjusted to bring ATTO647N fluorophores into the focal plane. Imager strand concentration was adjusted to ensure the single molecule condition, i.e. at most one imager strand binding event per MINFUX probing area, at an efficient binding frequency could be observed in confocal mode in the ROI. The entire ROI was then selected for MINFLUX imaging and imaged using the standard 2D MINFLUX imaging sequence with a hexagonal target-coordinate pattern (TCP), a 2% 640-nm-excitation power of 5.2 mW at periscope, and a pinhole size of 0.67 AU. Imager strand concentration was adjusted to obtain center frequency ratio (cfr, ratio between photon count detected during TCP center exposure and photon count detected during TCP offset exposures) values around < 0.7, with at least 40% of the localization attempts successfully passing the 4^th^ iteration, and effective frequency in offset (efo, background-corrected emission rates) values in the 80-100 kHz range. The imaging sequence was altered to include a localization limit (locLimit) of 10 per trace to allow for a more efficient sampling routine by avoiding oversampling of individual positions.

### MINFLUX data processing

MINFLUX data were filtered to remove traces with fewer than 3 localizations. The center of each trace was then found using Density-Based Spatial Clustering of Application with Noise (DBSCAN)^52^ with a search radius of 4 nm and a threshold of 3 localizations, yielding 2D coordinates for further analysis. Nearest neighbor analysis was then performed on these coordinates using the ball tree algorithm in Scikit-learn^53^. To identify isolated CD16a dimers, clusters of 2 trace centers in a 40-nm search radius were identified using DBSCAN and then filtered using the ball tree algorithm in Scikit-learn^53^. Analysis scripts can be found at https://github.com/pross1193/CD16-MINFLUX, and additional details on MINFLUX sample prep and data analysis can be found in our recently published book chapter^54^.

### Molecular dynamics modeling

As starting structure for our simulations, we used the cryo-EM structure of CD16a in complex with IgG Fc and both C21 and C28 nanobodies. Additionally, we performed simulations with CD16a solely in complex with C28, only in complex with the IgG Fc, only the Fc. For the complexes, we performed three repetitions of 1000 ns of classical molecular dynamics simulations using the AMBER 24 simulation software package which contains the pmemd.cuda module^55^. The structure was prepared using CHARMM-GUI^56–58^ and placed into cubic water boxes of TIP3P water molecules^59^ with a minimum wall distance to the protein of 12 Å and the charge was neutralized with K^+^ Cl^-^ ions up to a concentration of 0.15 mM. Parameters for the simulations were derived from the CHARMM36m force field^60^. The system was carefully equilibrated using a multistep equilibration protocol. Bonds involving hydrogen atoms were restrained using the SHAKE algorithm, allowing a timestep of 2.0 femtoseconds^61^. The Langevin thermostat was used to maintain the temperature during simulations at 300 K with a collision frequency of 2 ps^−1^ and a Monte Carlo barostat with one volume change attempt per 100 steps^62–64^.

For the cryo-EM complex, we calculated hydrogen bond and salt bridge interactions and frequencies using GetContacts (https://getcontacts.github.io/) and residue-wise interaction energies of the interacting residues using the LIE tool implemented in cpptraj^65^ to guide the design of the CD16a mutations. To quantify changes in flexibility, we calculated dihedral entropies using the X-entropy package^66^.

## Acknowledgements

We thank all patients and their families, as well as healthy blood donors and phlebotomists for participating in this research. NK-92/CD16a CRISPR cells were a kind gift from Dr. Steven Lin at Academia Sinica. NK-92/CD16a L48/H48 cells were a kind gift from Dr. Kerry Campbell at Fox Chase Cancer Center. YTS cells were a gift from Dr. Jordan Orange (University of Pennsylvania). With thank Dr. Nora Ross, currently at the Department of Applied Physics, Science for Life Laboratory, KTH Royal Institute of Technology, Stockholm, Sweden, for their thoughtful contribution to the content and interpretation of data in this study.

## Author contributions

T.C., H.F., B.C., and C.D.M. conducted NK cell culture, in vitro assays, and ADCC experiments; T.C., B.C., and C.D.M. prepared all proteins and reagents. E.R. and J.J. designed and performed LNP construction. C.D.M. led cryo-EM sample preparation, data collection, and structure determination, with protein contributions from T.C. and B.C. and computational support from M.F-Q. and J.L., who also performed molecular dynamics simulations. T.C. and C.D.M. performed MINFLUX sample preparation, data collection, and analysis with design and analytical contributions from J.M. and S.C.H.; STED and confocal imaging was carried out by T.C. and H.F. with experimental design and analysis support from K.S. D.P.L. prepared lipid nanoparticles and contributed to methods; R.L. and K.X. contributed to construct cloning and protein expression. M.B.Z., A.B.W., and E.M.M. provided experimental guidance and oversight. C.D.M. conceived and directed the study and drafted the manuscript, with writing and figure contributions from T.C., M.F-Q., H.F., J.L., E.R., J.M., S.C.H., K.S., and all senior authors.

## Data and materials availability

Cryo-EM density maps and atomic coordinates will be deposited in the Electron Microscopy Data Bank and Protein Data Bank upon manuscript submission but are also available upon request from the corresponding author. Molecular dynamics simulation trajectories and associated input files are available from the corresponding author upon reasonable request. All other data needed to evaluate the conclusions of this paper are present in the paper or the supplementary materials. Nanobodies C21 and C28, recombinant CD16a ectodomain variants, and LNP formulations described in this work are available from the corresponding author upon reasonable request, subject to a material transfer agreement.

## References

1. Mace, E. M. & Orange, J. S. New views of the human NK cell immunological synapse: recent advances enabled by super- and high-resolution imaging techniques. Front. Immunol. 3, 421 (2013).

2. Hazime, K. S. et al. Nanoscale restructuring of the immune synapse with an engager enhances NK cell function. Proc. Natl. Acad. Sci. 122, e2507336122 (2025).

3. Orange, J. S. Formation and function of the lytic NK-cell immunological synapse. Nat. Rev. Immunol. 8, 713–725 (2008).

4. Mace, E. M. et al. Cell biological steps and checkpoints in accessing NK cell cytotoxicity. Immunology and Cell Biology 92, 245–255 (2014).

5. Lou, Z., Jevremovic, D., Billadeau, D. D. & Leibson, P. J. A Balance between Positive and Negative Signals in Cytotoxic Lymphocytes Regulates the Polarization of Lipid Rafts during the Development of Cell-Mediated Killing. J. Exp. Med. 191, 347–354 (2000).

6. Galandrini, R. et al. SH2-containing inositol phosphatase (SHIP-1) transiently translocates to raft domains and modulates CD16-mediated cytotoxicity in human NK cells. Blood 100, 4581–4589 (2002).

7. Steblyanko, M., Anikeeva, N., Campbell, K. S., Keen, J. H. & Sykulev, Y. Integrins Influence the Size and Dynamics of Signaling Microclusters in a Pyk2-dependent Manner*. Journal of Biological Chemistry 290, 11833–11842 (2015).

8. March, M. E. & Long, E. O. β2 Integrin Induces TCRζ–Syk–Phospholipase C-γ Phosphorylation and Paxillin-Dependent Granule Polarization in Human NK Cells. J. Immunol. 186, 2998–3005 (2011).

9. Sugawara, S. et al. Knockdowns of CD3zeta Chain in Primary NK Cells Illustrate Modulation of Antibody-Dependent Cellular Cytotoxicity Against Human Immunodeficiency Virus-1. AIDS Res. Hum. Retroviruses 40, 631–636 (2024).

10. Phillips, J. H., Chang, C. & Lanier, L. L. Platelet-induced expression of FcγRIII (CD16) on human monocytes. Eur. J. Immunol. 21, 895–899 (1991).

11. Zhou, Q., Gil-Krzewska, A., Peruzzi, G. & Borrego, F. Matrix metalloproteinases inhibition promotes the polyfunctionality of human natural killer cells in therapeutic antibody-based anti-tumour immunotherapy. Clin. Exp. Immunol. 173, 131–139 (2013).

12. Ma, R., Garcia, K. C., Cui, B. & Covert, M. W. Antibody Fc receptor CD16a mediates natural killer cell activation via mechanotransduction of piconewton forces. (2026) doi:10.64898/2026.02.06.704366.

13. Ross, P. et al. CD16a pairs form the basal molecular subunit for the NK-cell ADCC lytic synapse. J. Immunol. (2025) doi:10.1093/jimmun/vkaf077.

14. Kremer, P. G. et al. The impact of N-glycan conformational entropy on the binding affinity of Fc γ receptor IIIa/CD16a. Structure 34, 454–462.e4 (2026).

15. Perussia, B., Starr, S., Abraham, S., Fanning, V. & Trinchieri, G. Human natural killer cells analyzed by B73.1, a monoclonal antibody blocking Fc receptor functions. I. Characterization of the lymphocyte subset reactive with B73.1. J. Immunol. (Baltim., Md : 1950) 130, 2133–41 (1983).

16. Grier, J. T. et al. Human immunodeficiency-causing mutation defines CD16 in spontaneous NK cell cytotoxicity. J. Clin. Investig. 122, 3769–3780 (2012).

17. Monteiro, M. F. et al. NK Cytotoxicity Mediated by NK-92 Cell Lines Expressing Combinations of Two Allelic Variants for FCGR3. Antibodies 13, 55 (2024).

18. Maskalenko, N. A. et al. The FcγRIIIA (CD16) L48-H/R polymorphism enhances NK cell-mediated antibody-dependent cellular cytotoxicity by promoting serial killing. Cancer Immunol. Res. (2024) doi:10.1158/2326-6066.cir-24-0384.

19. Behar, G. et al. Isolation and characterization of anti-FcγRIII (CD16) llama single-domain antibodies that activate natural killer cells. Protein Eng., Des. Sel. 21, 1–10 (2008).

20. Chen, M., Su, Q. & Shi, Y. Molecular mechanism of IgE-mediated FcεRI activation. Nature 637, 453–460 (2025).

21. Huang, R.-S., Shih, H.-A., Lai, M.-C., Chang, Y.-J. & Lin, S. Enhanced NK-92 Cytotoxicity by CRISPR Genome Engineering Using Cas9 Ribonucleoproteins. Front Immunol 11, 1008 (2020).

22. Maskalenko, N. A. et al. The FcγRIIIA (CD16) L48-H/R polymorphism enhances NK cell-mediated antibody-dependent cellular cytotoxicity by promoting serial killing. Cancer Immunol. Res. (2024) doi:10.1158/2326-6066.cir-24-0384.

23. Benavente, M. C. R., Hughes, H. B., Kremer, P. G., Subedi, G. P. & Barb, A. W. Inhibiting N-glycan processing increases the antibody binding affinity and effector function of human natural killer cells. Immunology 170, 202–213 (2023).

24. Benavente, M. C. R. et al. Distinct CD16a features on human NK cells observed by flow cytometry correlate with increased ADCC. Sci. Rep. 14, 7938 (2024).

25. Kakiuchi-Kiyota, S. et al. A BCMA/CD16A bispecific innate cell engager for the treatment of multiple myeloma. Leukemia 36, 1006–1014 (2022).

26. Ahmed, A. A., Keremane, S. R., Vielmetter, J. & Bjorkman, P. J. Structural characterization of GASDALIE Fc bound to the activating Fc receptor FcγRIIIa. J Struct Biol 194, 78–89 (2016).

27. Roberts, J. T., Patel, K. R. & Barb, A. W. Site-specific N-glycan Analysis of Antibody-binding Fc γ Receptors from Primary Human Monocytes*. Mol. Cell. Proteom. 19, 362–374 (2020).

28. Blázquez-Moreno, A., et al. Transmembrane features governing Fc receptor CD16A assembly with CD16A signaling adaptor molecules. Proc. Natl. Acad. Sci. 114, E5645–E5654 (2017).

29. Lanier, L. L., Yu, G. & Phillips, J. H. Co-association of CD3ζ with a receptor (CD16) for IgG Fc on human natural killer cells. Nature 342, 803–805 (1989).

30. Anderson, P. et al. Fc gamma receptor type III (CD16) is included in the zeta NK receptor complex expressed by human natural killer cells. Proc. Natl. Acad. Sci. 87, 2274–2278 (1990).

31. Delcassian, D. et al. Nanoscale Ligand Spacing Influences Receptor Triggering in T Cells and NK Cells. Nano Lett 13, 5608–5614 (2013).

32. Vivier, E. et al. Signaling function of reconstituted CD16: ζ:γ receptor complex isoforms. Int. Immunol. 4, 1313–1323 (1992).

33. Strauss, S. & Jungmann, R. Up to 100-fold speed-up and multiplexing in optimized DNA-PAINT. Nat. Methods 17, 789–791 (2020).

34. Jungmann, R. et al. Multiplexed 3D cellular super-resolution imaging with DNA-PAINT and Exchange-PAINT. Nat. Methods 11, 313–318 (2014).

35. Mukhortava, A. & Schlierf, M. Efficient Formation of Site-Specific Protein–DNA Hybrids Using Copper-Free Click Chemistry. Bioconjugate Chem. 27, 1559–1563 (2016).

36. Delehedde, C. et al. Enhancing natural killer cells proliferation and cytotoxicity using imidazole-based lipid nanoparticles encapsulating interleukin-2 mRNA. Mol. Ther. - Nucleic Acids 35, 102263 (2024).

37. Douka, S. et al. Lipid nanoparticle-mediated messenger RNA delivery for ex vivo engineering of natural killer cells. J. Control. Release 361, 455–469 (2023).

38. Patel, S. et al. Naturally-occurring cholesterol analogues in lipid nanoparticles induce polymorphic shape and enhance intracellular delivery of mRNA. Nat. Commun. 11, 983 (2020).

39. Robinson, J. I., et al. Affimer proteins inhibit immune complex binding to FcγRIIIa with high specificity through competitive and allosteric modes of action. Proc. Natl. Acad. Sci. 115, E72–E81 (2018).

40. Kondadasula, S. V. et al. Colocalization of the IL-12 receptor and FcγRIIIa to natural killer cell lipid rafts leads to activation of ERK and enhanced production of interferon-γ. Blood 111, 4173–4183 (2008).

41. Kara, S. et al. Impact of Plasma Membrane Domains on IgG Fc Receptor Function. Front. Immunol. 11, 1320 (2020).

42. Bryan, A. M. et al. Cholesterol and sphingomyelin are critical for Fcγ receptor–mediated phagocytosis of Cryptococcus neoformans by macrophages. J. Biol. Chem. 297, 101411 (2021).

43. Chen, J. et al. Tuning charge density of chimeric antigen receptor optimizes tonic signaling and CAR-T cell fitness. Cell Res. 33, 341–354 (2023).

44. Binyamin, L. et al. Blocking NK Cell Inhibitory Self-Recognition Promotes Antibody-Dependent Cellular Cytotoxicity in a Model of Anti-Lymphoma Therapy. J Immunol 180, 6392–6401 (2008).

45. Miah, S. M. S. & Campbell, K. S. Natural Killer Cell Protocols, Cellular and Molecular Methods. Methods Mol. Biol. 612, 199–208 (2009).

46. Subedi, G. P., Johnson, R. W., Moniz, H. A., Moremen, K. W. & Barb, A. High Yield Expression of Recombinant Human Proteins with the Transient Transfection of HEK293 Cells in Suspension. J. Vis. Exp. e53568 (2015) doi:10.3791/53568.

47. Pettersen, E. F. et al. UCSF Chimera—A visualization system for exploratory research and analysis. J. Comput. Chem. 25, 1605–1612 (2004).

48. Meng, E. C. et al. UCSF ChimeraX: Tools for structure building and analysis. Protein Sci. 32, e4792 (2023).

49. Salentin, S., Schreiber, S., Haupt, V. J., Adasme, M. F. & Schroeder, M. PLIP: fully automated protein–ligand interaction profiler. Nucleic Acids Res. 43, W443–W447 (2015).

50. Vlatkovic, I. et al. Ribozyme Assays to Quantify the Capping Efficiency of In Vitro-Transcribed mRNA. Pharmaceutics 14, 328 (2022).

51. Schmidt, R. et al. MINFLUX nanometer-scale 3D imaging and microsecond-range tracking on a common fluorescence microscope. Nat. Commun. 12, 1478 (2021).

52. Ester, M., Kriegel, H.-P., Sander, J. & Xu, X. A density-based algorithm for discovering clusters in large spatial databases with noise. kdd 226–231 (1996).

53. Pedregosa et al. Scikit-learn: Machine Learning in Python. The Journal of Machine Learning Research 12, 2825–2830 (2011).

54. Ross, N. et al. MINFLUX nanoscopy to study the NK cell immune synapse. Methods Cell Biol. (2026) doi:10.1016/bs.mcb.2026.01.021.

55. Case, D. A. et al. Recent Developments in Amber Biomolecular Simulations. J. Chem. Inf. Model. 65, 7835–7843 (2025).

56. Lee, J. et al. CHARMM-GUI Input Generator for NAMD, GROMACS, AMBER, OpenMM, and CHARMM/OpenMM Simulations Using the CHARMM36 Additive Force Field. J. Chem. Theory Comput. 12, 405–413 (2016).

57. Lee, J. et al. CHARMM-GUI supports the Amber force fields. J. Chem. Phys. 153, 035103 (2020).

58. Jo, S., Kim, T., Iyer, V. G. & Im, W. CHARMM-GUI: A web-based graphical user interface for CHARMM. J. Comput. Chem. 29, 1859–1865 (2008).

59. Jorgensen, W. L., Chandrasekhar, J., Madura, J. D., Impey, R. W. & Klein, M. L. Comparison of simple potential functions for simulating liquid water. J. Chem. Phys. 79, 926–935 (1983).

60. Huang, J. et al. CHARMM36m: an improved force field for folded and intrinsically disordered proteins. Nat. Methods 14, 71–73 (2017).

61. Andersen, H. C. Rattle: A “velocity” version of the shake algorithm for molecular dynamics calculations. J. Comput. Phys. 52, 24–34 (1983).

62. Åqvist, J., Wennerström, P., Nervall, M., Bjelic, S. & Brandsdal, B. O. Molecular dynamics simulations of water and biomolecules with a Monte Carlo constant pressure algorithm. Chem. Phys. Lett. 384, 288–294 (2004).

63. Doll, J. D., Myers, L. E. & Adelman, S. A. Generalized Langevin equation approach for atom/solid-surface scattering: Inelastic studies. J. Chem. Phys. 63, 4908–4914 (1975).

64. Adelman, S. A. & Doll, J. D. Generalized Langevin equation approach for atom/solid-surface scattering: General formulation for classical scattering off harmonic solids. J. Chem. Phys. 64, 2375–2388 (1976).

65. Roe, D. R. & Cheatham, T. E. PTRAJ and CPPTRAJ: Software for Processing and Analysis of Molecular Dynamics Trajectory Data. J. Chem. Theory Comput. 9, 3084–3095 (2013).

66. Kraml, J., Hofer, F., Quoika, P. K., Kamenik, A. S. & Liedl, K. R. X-Entropy: A Parallelized Kernel Density Estimator with Automated Bandwidth Selection to Calculate Entropy. J. Chem. Inf. Model. 61, 1533–1538 (2021).

67. Lord, S. J., Velle, K. B., Mullins, R. D. & Fritz-Laylin, L. K. SuperPlots: Communicating reproducibility and variability in cell biology. J. Cell Biol. 219, e202001064 (2020).

