## Supplementary Materials for "The membrane distal domain of CD16a allosterically regulates NK cell ADCC"

Tania Cid *et al.*

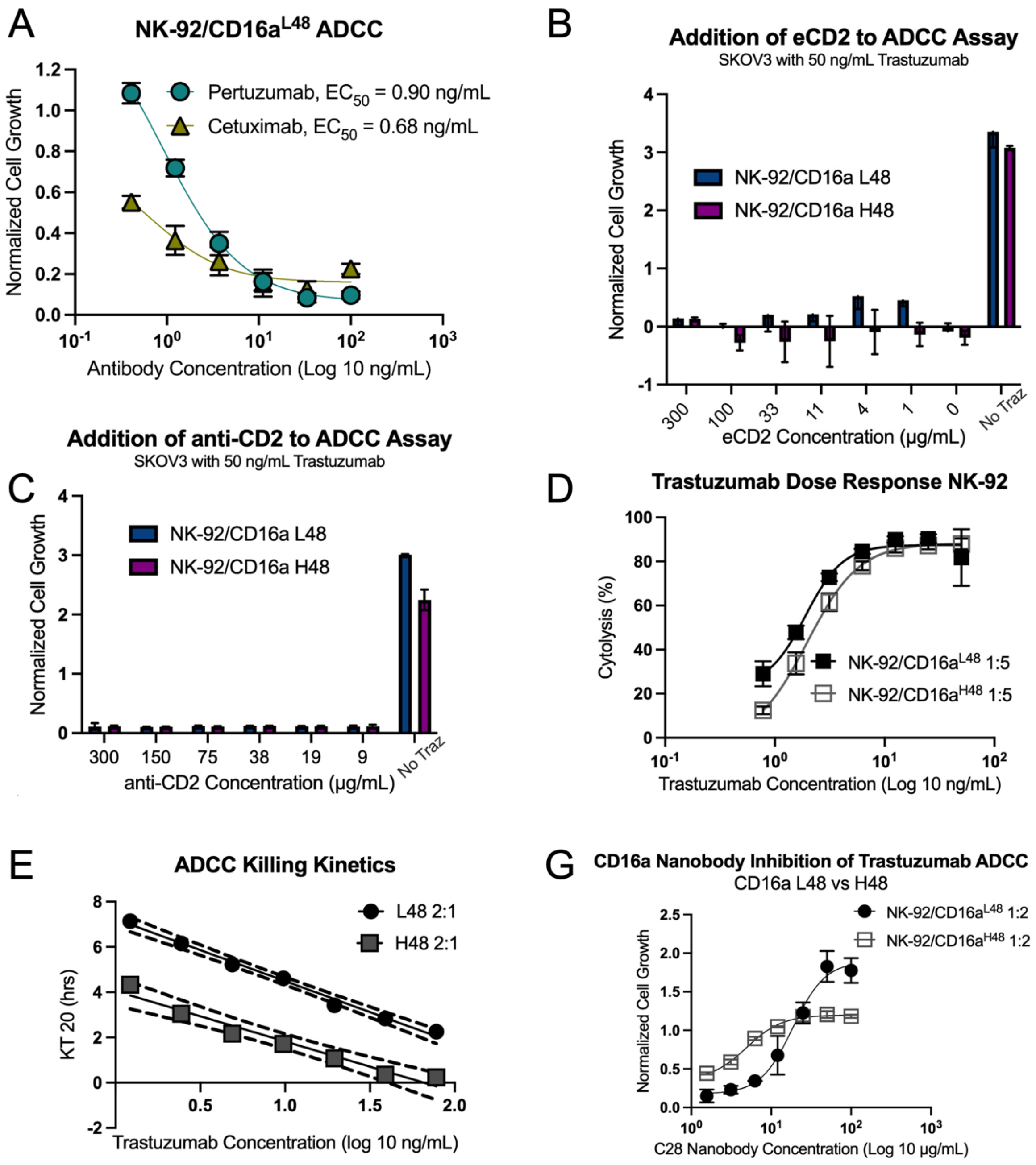


**Figure S1. Additional NK cell cytotoxicity data.** A) xCELLigence real-time cell analysis (RTCA) ADCC assay, using SKOV3 target cells, coated in Pertuzumab or Cetuximab, and NK-92/CD16a$^{L48}$ cells as effector cells at an E:T of 2:1. Normalized cell growth corresponds to target cell death. Endpoint data from 14h post-assay start. B) The extracellular domain of CD2 (eCD2) was added at the indicated concentrations to SKOV3 cells coated in Trastuzumab. NK-92/CD16a cells were added at a ratio of 2:1 and normalized cell growth at 14 hours post-assay start is graphed. C) Identical assay as in B but cells were instead coated in an anti-human CD2 antibody at the indicated concentrations. D) RTCA ADCC assay as in part A but comparing NK-92 cell lines generated using lentiviruses packaging CD16a$^{L48}$ or CD16a$^{H48}$ at an E:T ratio of 1:5. E) Killing kinetics of assay from part D at an E:T ratio of 2:1. Dotted lines represent standard deviation from the mean. Plotted are the kill time to 20% cytotoxicity (KT20) at each indicated concentration of Trastuzumab. G) C28-inhibition of NK-92/CD16a ADCC activity, using Trastuzumab coated SKOV3 cells (100 ng/mL) and increasing concentrations of C28 nanobody at an E:T of 2:1. As nanobody concentration increases, normalized cell growth is restored even at a maximally inhibitory concentration of Trastuzumab. $EC_{50}$ is the effective concentration at 50% cytotoxicity. $IC_{50}$ is the inhibitory concentration at 50% antibody inhibition.

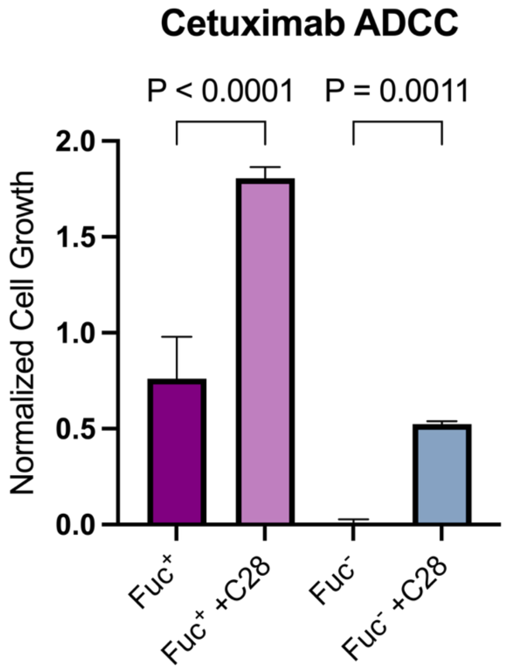


**Figure S2. ADCC activity of fucose versus fucose-free Cetuximab with and without C28 nanobody.** Cetuximab was produced in CHO cells without (Fuc$^{+}$) and with (Fuc$^{2}$) a fucose inhibitor and then used to test ADCC activity in the absence or presence of C28. Cetuximab is at 100 ng/mL and C28 is at 7.5 µg/mL. IgG containing fucose results in a 2.4-fold difference in activity with C28 present, whereas afucosylated IgG results in a 63-fold difference. Normalized cell growth was assessed at 12 hours post NK cell addition. Statistical significance was determined using an ordinary one-way ANOVA test (n = 3).

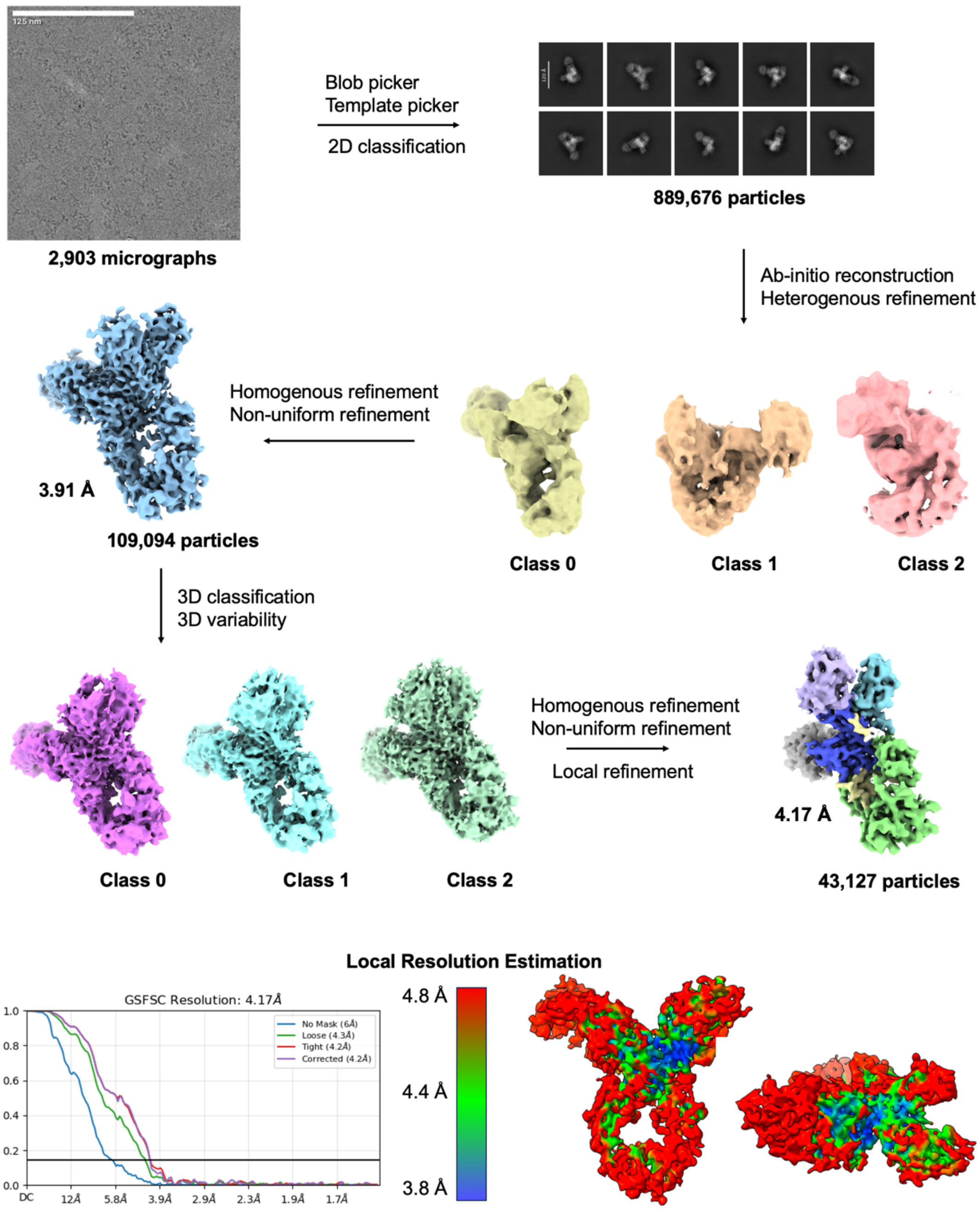


**Figure S3. Cryo-EM data processing workflow.**

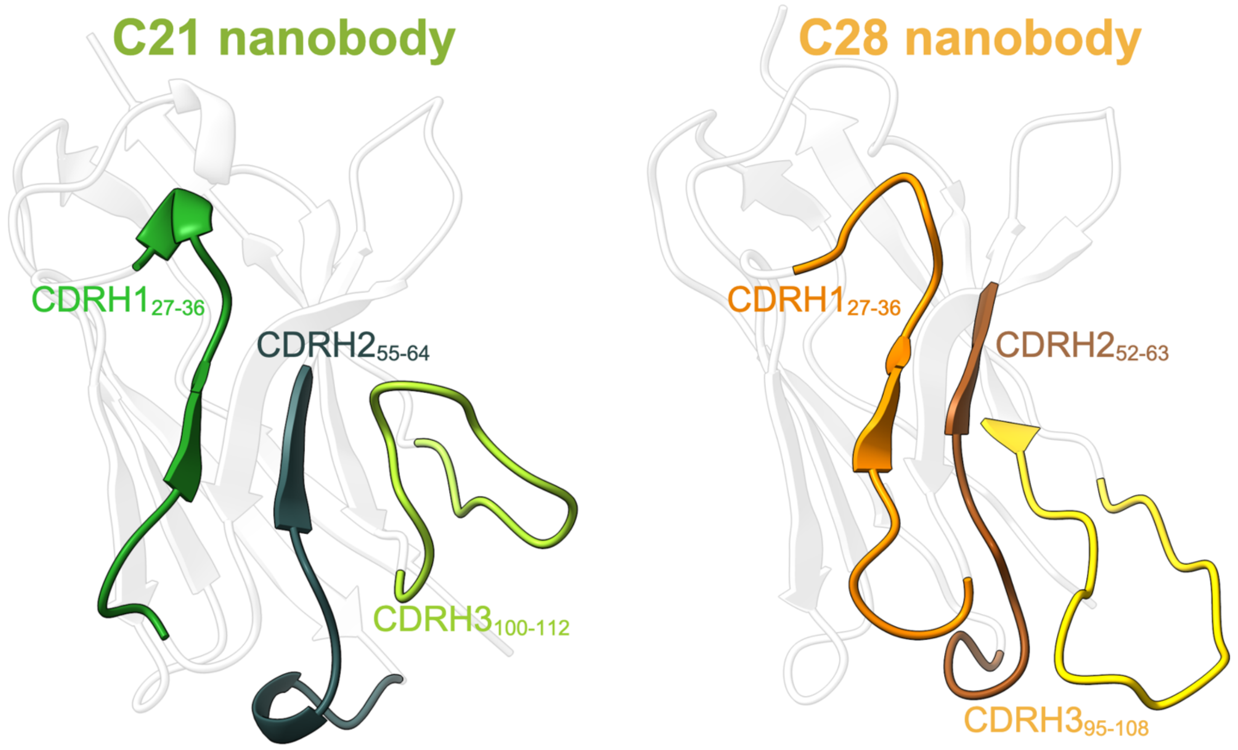


**Figure S4. Paratope details of C21 and C28 nanobodies.** Models of nanobodies C21 and C28 were built into the final cryo-EM map (Fig. 3) and the complementary determining regions (CDRs) are indicated here. CDR assignments were made according to the IMGT format, numbering sequentially from the first amino acid after the signal peptide.

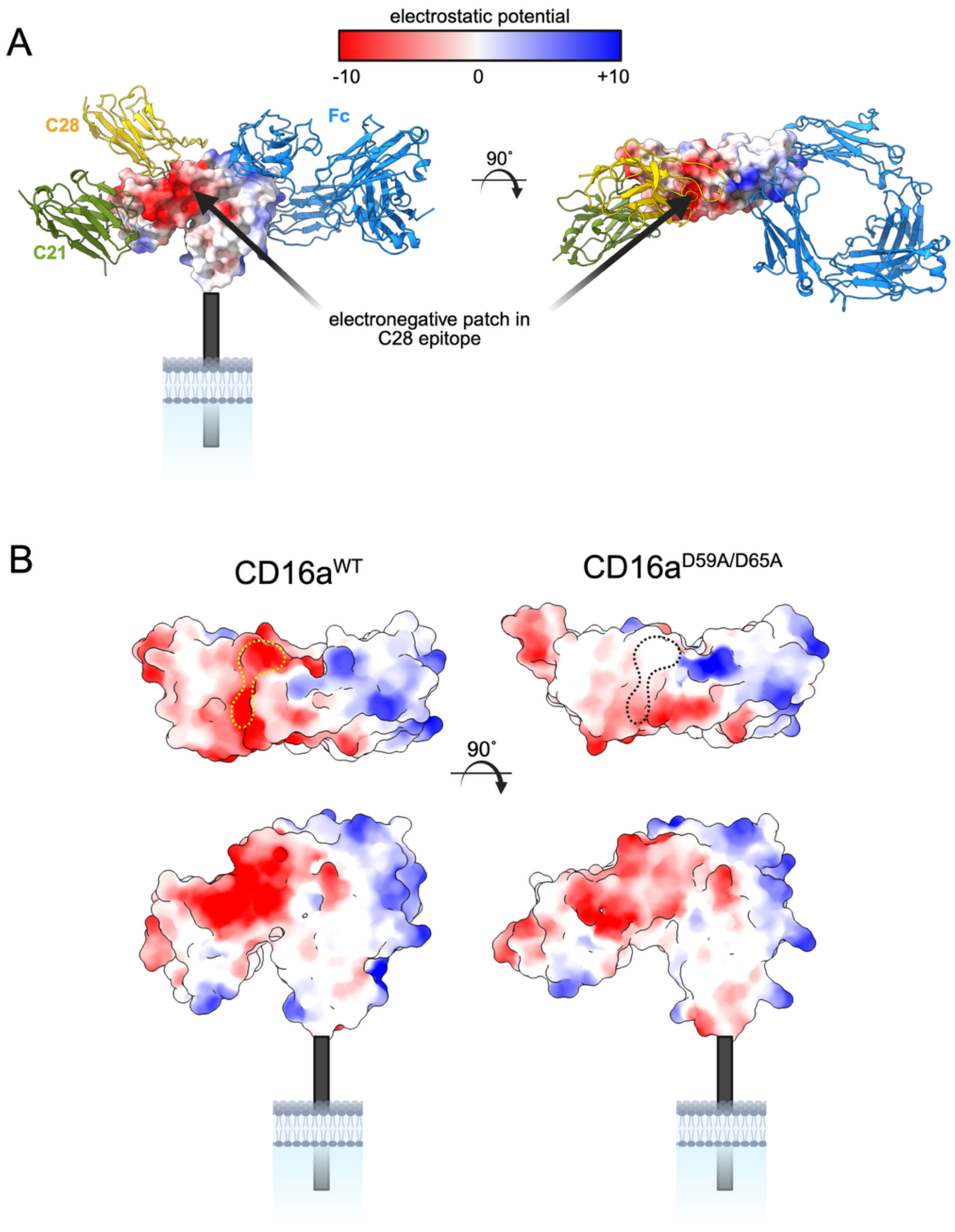


**Figure S5. Electrostatic potential of CD16a in relation to the C21 and C28 epitopes.** A) Mapping the electrostatic surface potential onto our modeled CD16a structure, we note that C28 overlaps with a large, highly electronegative patch (red) that extends from the top to the side of D1. B) Comparing our CD16a D59A/D65A mutant to WT, we see that the electronegative patch comprising the C28 epitope shifts to neutral in charge (yellow dotted lines and black dotted lines for WT and D59A/D65A, respectively).

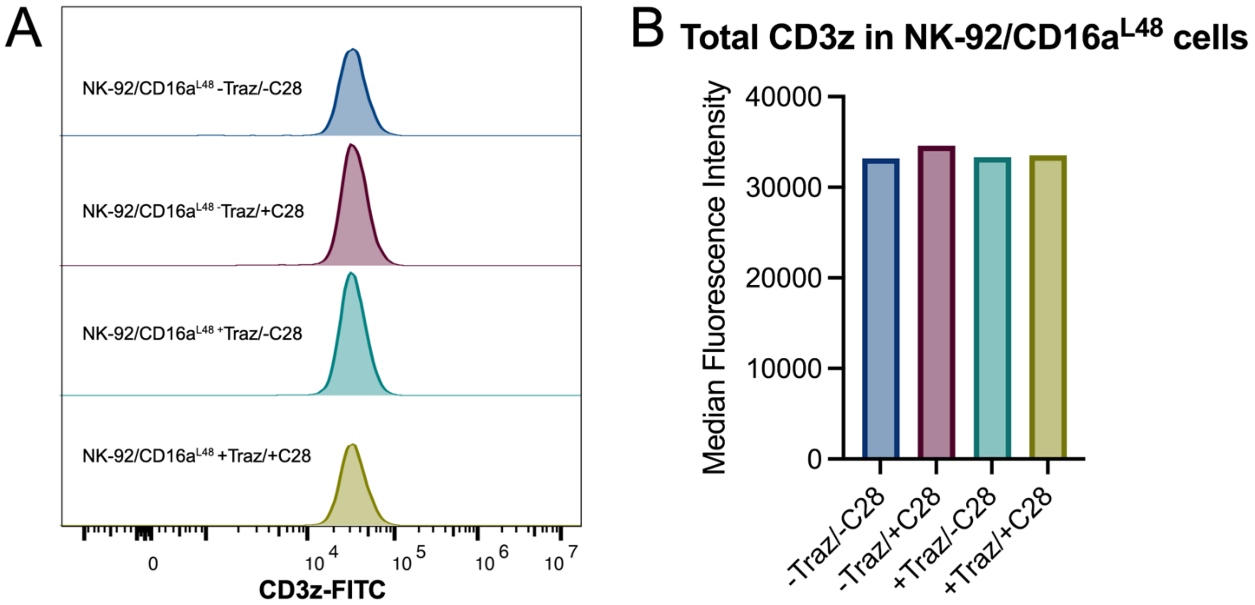


**Figure S6. Total CD3z does not change upon NK cell activation on SLBs, with or without C28.** A) NK-92/CD16a$^{L48}$ cells were activated on SLBs containing ICAM/HER2 with or without Trastuzumab and with or without C28. After activation, cells were harvested, fixed and permeabilized, stained for total CD3z, and analyzed by flow cytometry. B) No noticeable differences in total CD3z, as measured by median fluorescence intensity, were observed under any of the conditions.

A

**LNP characterization**

| | EE (%) | size (nm) | PDI |
|---|---|---|---|
| chLNP | 91 | 106 ± 38 | 0.133 |
| ch5LNP | 95 | 82 ± 26 | 0.104 |
| bLNP | 76 | 148 ± 60 | 0.169 |
| b5LNP | 94 | 121 ± 53 | 0.192 |

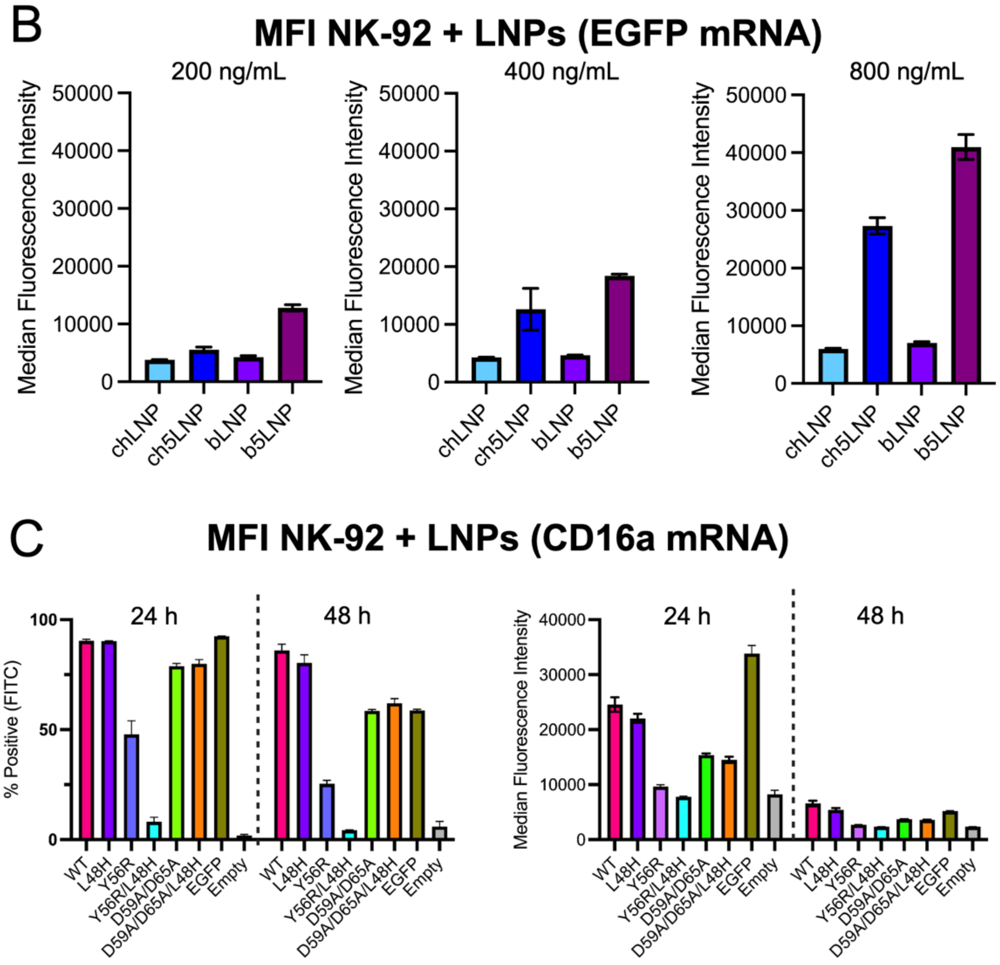


**Figure S7. Validation of lipid nanoparticle (LNP) target gene protein expression in NK-92 cells.** A) Encapsulation efficiency (EE) determined by Ribogreen prior to addition of cryoprotectant. Z-avg diameter (size) and polydispersity index (PDI) are average of n = 8 multi-angle light scattering readings ± SD where applicable. B) LNPs at the indicated formulation and containing the amount of EGFP mRNA indicated were used to transfect NK-92 cells. After 24h, cells were checked for A) positive expression of EGFP (using FITC) and B) median fluorescence intensity (MFI). C) The number of positive cells (left) and MFI (right) of CD16a constructs were measured by flow cytometry at 24 and 48h post-transfection, using 400 ng/mL b5LNPs with 5%

(w/v) sucrose. CD16a expression was measured using anti-human CD16a (3G8) conjugated to Alexa Fluor-488. bLNPs refer to LNPs with beta-sitosterol and chLNPs refer to LNPs with cholesterol, both with an nucleic acid (N) to phospholipid (P) ratio of 1:1. b5LNPs and ch5LNPs have an N:P ratio of 5:1.
